# MTOR modulation induces selective perturbations in histone methylation which influence the anti-proliferative effects of mTOR inhibitors

**DOI:** 10.1101/2023.11.07.566067

**Authors:** HaEun Kim, Benjamin Lebeau, David Papadopoli, Predrag Jovanovic, Mariana Russo, Daina Avizonis, Masahiro Morita, Farzaneh Afzali, Josie Ursini-Siegel, Lynne-Marie Postovit, Michael Witcher, Ivan Topisirovic

## Abstract

Emerging data suggest a significant cross-talk between metabolic and epigenetic programs. However, the relationship between the mechanistic target of rapamycin (mTOR) which is a pivotal regulator of cellular metabolism, and epigenetic modifications remains poorly understood. We thus explored the impact of modulating mTOR signaling on histone methylation, a well-known epigenetic modification. Our results showed that mTORC1 activation caused by abrogation of TSC2 increased H3K27me3 but not H3K4me3 or H3K9me3. This appeared to be mediated via the induction of EZH2 protein synthesis, downstream of 4EBPs. Surprisingly, mTOR inhibition also induced H3K27me3 independently of TSC2. This coincided with reduced EZH2 and increased EZH1 protein levels. Notably, the ability of mTOR inhibitors to induce H3K27me3 levels was positively correlated with their anti-proliferative effects. Collectively, our findings demonstrate that both activation and inhibition of mTOR selectively increase H3K27me3 by distinct mechanisms, whereby the ability of mTOR inhibitors to induce H3K27me3 influences their anti-proliferative effects.

**Highlights:**

- Paradoxically, both mTOR activation and inhibition induce H3K27me3.
- The effect of mTOR inhibitors on H3K27me3 are not secondary to cell cycle arrest.
- H3K27me3 triggered by mTOR suppression coincides with perturbations in EZH1/2 ratio.
- H3K27me3 impacts on the anti-proliferative effects of mTOR inhibitors.

## Introduction

The mechanistic target of rapamycin (mTOR) signaling pathway plays a pivotal role in adjusting metabolic programs in response to a variety of extracellular stimuli and intracellular cues in both homeostatic and pathological contexts ^1,2^. In mammals, mTOR is involved in two distinct complexes: mTOR complex 1 (mTORC1) and mTOR complex 2 (mTORC2) that differ in their composition and function ^1,2^. mTORC1 stimulates protein synthesis, various metabolic pathways, and suppresses autophagy to bolster cellular growth and proliferation ^1–3^. In turn, mTORC2 governs cytoskeletal organization, survival, and glucose and lipid metabolism, whereby the effects of mTORC2 on metabolism are largely distinct from those of mTORC1 ^4–6^. The effects of mTORC1 and mTORC2 are also mediated by distinct substrates. The best characterized mTORC1 substrates to date are the ribosomal protein S6 kinases (S6Ks, 1 and 2 in mammals), and eukaryotic translation initiation factor 4E (eIF4E)-binding proteins (4E-BPs; 1-3 in mammals), which mediate their effects on protein synthesis ^1,2^. Known mTORC2 substrates comprise several AGC kinase family members including AKT and serum- and glucocorticoid-inducible kinase 1 (SGK1) ^7,8^. Finally, mTORC1 and mTORC2 differ in their sensitivity to the allosteric inhibitor rapamycin, which directly inhibits mTORC1 but not mTORC2 ^9^. In addition to its role in homeostasis, dysregulation of mTOR signaling plays a prominent role across human pathologies including neoplasia ^10,11^. Indeed, mTOR signaling is frequently upregulated in a broad range of cancers due to oncogene activation (e.g., *PIK3CA* and *RAS*) or inactivation of tumor suppressors (e.g., *PTEN*, *TSC1/2*) that impinge on the mTOR pathway ^12–15^. This renders mTOR an appealing target for the development of anti-cancer treatments ^16^.

Epigenetic modifications are inheritable and potentially reversible alterations that modulate gene expression without altering the DNA sequence ^17^. Similar to dysregulation of mTOR signaling, disruption of epigenetic regulation is thought to play a prominent role in neoplasia ^18^. Accordingly, widespread alterations in the epigenetic landscape are frequently observed in cancer ^19^. One of the most prominent epigenetic modifications that has been linked to cancer is histone methylation, which is a multifaceted and dynamic process occurring at various locations on the histone tail ^20^. Histone methylation has diverse effects on gene expression ^20^. For instance, the trimethylation of lysine 4 on the histone H3 subunit (H3K4me3) is commonly linked with transcriptional activation, whereas H3K9me3 and H3K27me3 are associated with the suppression of transcription ^21^.

While the involvement of mTOR in various biological processes including protein synthesis, metabolic regulation, and autophagy has been extensively explored ^22^, its role in epigenetic reprogramming remains largely uncharted. Recent studies have indicated that mTOR may regulate histone methylation by modulating levels and/or activity of epigenetic modifiers such as Enhancer of zeste homolog 2 (EZH2), Jumonji domain containing 1C (JMJD1C), and G9a ^23–25^. Also, considering its prominent role in governing glucose metabolism and the serine-glycine-one-carbon pathway ^26–28^, mTOR may influence epigenetic modifications indirectly, by modulating levels of methyl donors [e.g., S-Adenosyl methionine (SAM)] or metabolites that influence the function of histone demethylases [e.g., α-ketoglutarate (α-KG), fumarate, and succinate] ^29,30^.

Notwithstanding these potential links between mTOR signaling and epigenetic programs, there is a general paucity of studies exploring potential connections between mTOR and epigenetic modifications. To address this, we focused on three well-characterized histone methylation marks H3K4me3, H3K9me3 and H3K27me3. Surprisingly, we observed that both hyperactivation and inhibition of mTOR selectively increased H3K27me3, but not H3K4me3 and H3K9me3 levels. mTORC1 hyperactivation was paralleled by the induction of H3K27me3 via a 4E-BP-dependent increase in EZH2 protein levels. In contrast, induction of H3K27me3 upon mTOR inhibition was accompanied by increase in EZH1 and decrease in EZH2 levels. Finally, we provide evidence that H3K27me3 is at least in part required for the anti-proliferative effects of mTOR inhibitors. Collectively, these findings establish a hitherto unappreciated link between mTOR and histone methylation and suggest that epigenetic mechanisms may partially be responsible for the anti-proliferative effects of mTOR inhibitors.

## Results

### Hyperactivation of mTORC1 drives increase in H3K27me3 via the 4E-BPs/EZH2 axis

To investigate the impact of mTORC1 hyperactivation on histone methylation, we first employed *TSC2*-null mouse embryonic fibroblasts (MEFs) ^31^. TSC2 functions as a negative regulator of the mTOR pathway by acting as a GTPase-activating protein (GAP) for Ras homolog enriched in brain (Rheb) and thus suppressing Rheb-dependent activation of mTORC1 ^32^. As anticipated, relative to wild-type (WT) *TSC2*-proficient MEFs, *TSC2*-null MEFs demonstrated higher phosphorylation levels of mTORC1 substrate 4E-BP1 (S65) and ribosomal protein S6 [rpS6 (S240/244)] that is phosphorylated via the mTORC1/S6K axis ^3^ (Figure 1A). To examine the effects of TSC2 loss and subsequent activation of mTORC1 on histone methylation, we compared the levels of three methylation marks (H3K4me3, H3K9me3, and H3K27me3) between *TSC2* WT and KO cells. These experiments revealed that H3K27me3 is higher in *TSC2*-null cells relative to WT cells (Figure 1A). In contrast, the levels of H3K4me3 and H3K9me3 remained largely unaffected by the TSC2 status and mTORC1 activity in the cell (Figure 1A). These findings suggest that increased mTORC1 signaling caused by TSC2 loss is paralleled by selective perturbations in histone methylation, specifically an increase in H3K27me3, while H3K4me3 and H3K9me3 remain unchanged.

**Figure 1.**
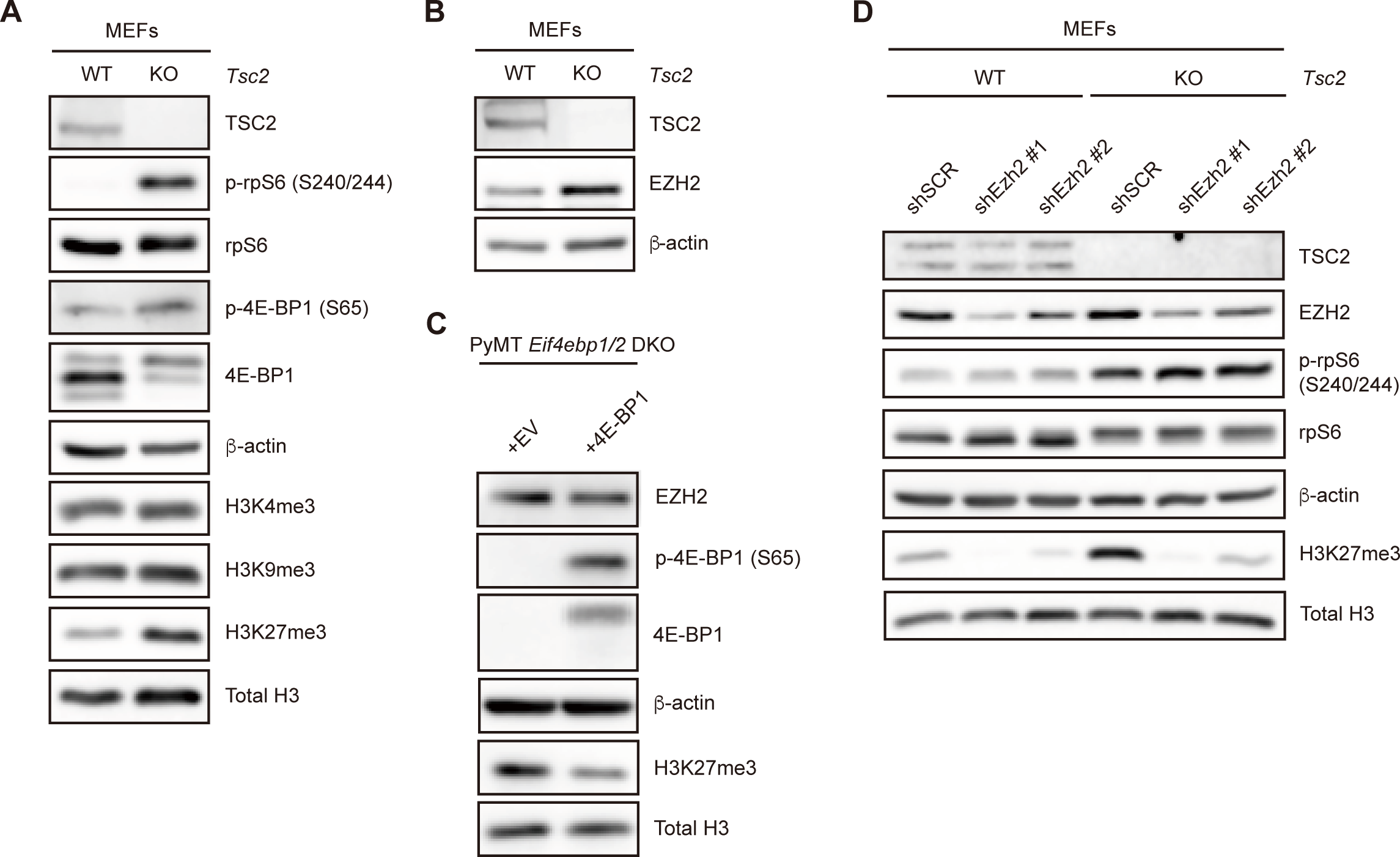
Constitutive mTORC1 activation selectively induces H3K27me3 via the 4E-BPs/EZH2 axis. (A) Levels of the indicated proteins in *TSC2* WT and KO MEFs were determined by Western blotting (n=3). β-actin served as a loading control. (B) Levels of the indicated proteins in *TSC2* WT and KO MEFs were determined by Western blotting (n=3). β-actin served as a loading control. (C) Levels of the indicated proteins in *Eif4ebp1/2* DKO cells infected with EV or human 4E-BP1 were assessed by Western blotting (n=3). β-actin served as a loading control. (D) Levels of the indicated proteins in *TSC2* WT and KO MEFs infected with a scrambled (shSCR) or Ezh2-specific shRNA (shEzh2) were determined by Western blotting (n=3). β-actin served as a loading control.

EZH2 is a component of the polycomb repressive complex 2 (PRC2) that catalyzes H3K27 methylation ^33^. It has been previously demonstrated that EZH2 protein synthesis is elevated in breast cancer cells where increased mTORC1 activity is driven by c-SRC ^23^. To test the generality of these findings, we investigated the impact of mTORC1 activation caused by TSC2 loss on EZH2 protein levels. Consistent with the c-SRC-driven mTORC1 hyperactivation, *TSC2*-null MEFs displayed higher levels of EZH2 protein in comparison to their WT counterparts (Figure 1B). EZH2 protein synthesis is increased via the mTORC1/4E-BPs axis in the context of c-SRC activation ^23^. Indeed, using breast cancer cells isolated from MMTV-PyMT/*Eif4ebp1/2*-null mice, we observed that re-expression of 4E-BP1 coincides with a decrease in EZH2 protein levels and downregulation of H3K27me3 (Figure 1C). In summary, these results show that mTORC1 regulates H3K27me3 in a 4E-BPs-dependent manner under various contexts, including c-SRC activation ^23^ and TSC2 loss. We next set out to confirm that the alterations in H3K27me3 levels observed in *TSC2* KO MEFs were indeed mediated by EZH2. To address this, we depleted EZH2 in *TSC2* WT or KO MEFs using shRNAs. EZH2 depletion abolished H3K27me3 levels in both *TSC2* WT and KO MEFs (Figure 1D). Overall, these findings demonstrate that mTORC1 activation coincides with the induction of H3K27me3 which is mediated via the 4E-BPs/EZH2 axis.

### Inhibition of mTOR is paralleled by selective perturbations in histone methylation

After establishing the effects of mTORC1 activation on histone methylation, we next sought to investigate whether mTOR inhibition will reduce H3K27me3 levels in TSC2 proficient or deficient MEFs. In these experiments, we subjected *TSC2* WT or KO MEFs to overnight serum starvation, upon which cells were stimulated by 10% fetal bovine serum (FBS) in the presence of active-site mTOR inhibitor Ink128 or a vehicle (DMSO) for 4, 24 or 48 hours. While acute mTOR inhibition (4 hours) had no appreciable effect on H3K27me3 levels, prolonged mTOR inhibition (24 and 48 hours), surprisingly induced H3K27me3 levels in both *TSC2* WT and KO MEFs (Figure 2A). Importantly, Ink128-induced elevation in H3K27me3 levels occurred independently of the TSC2 status in the cell (Figure 2A). These findings suggest that paradoxically both mTORC1 activation and mTOR inhibition result in upregulation of H3K27me3 levels.

**Figure 2.**
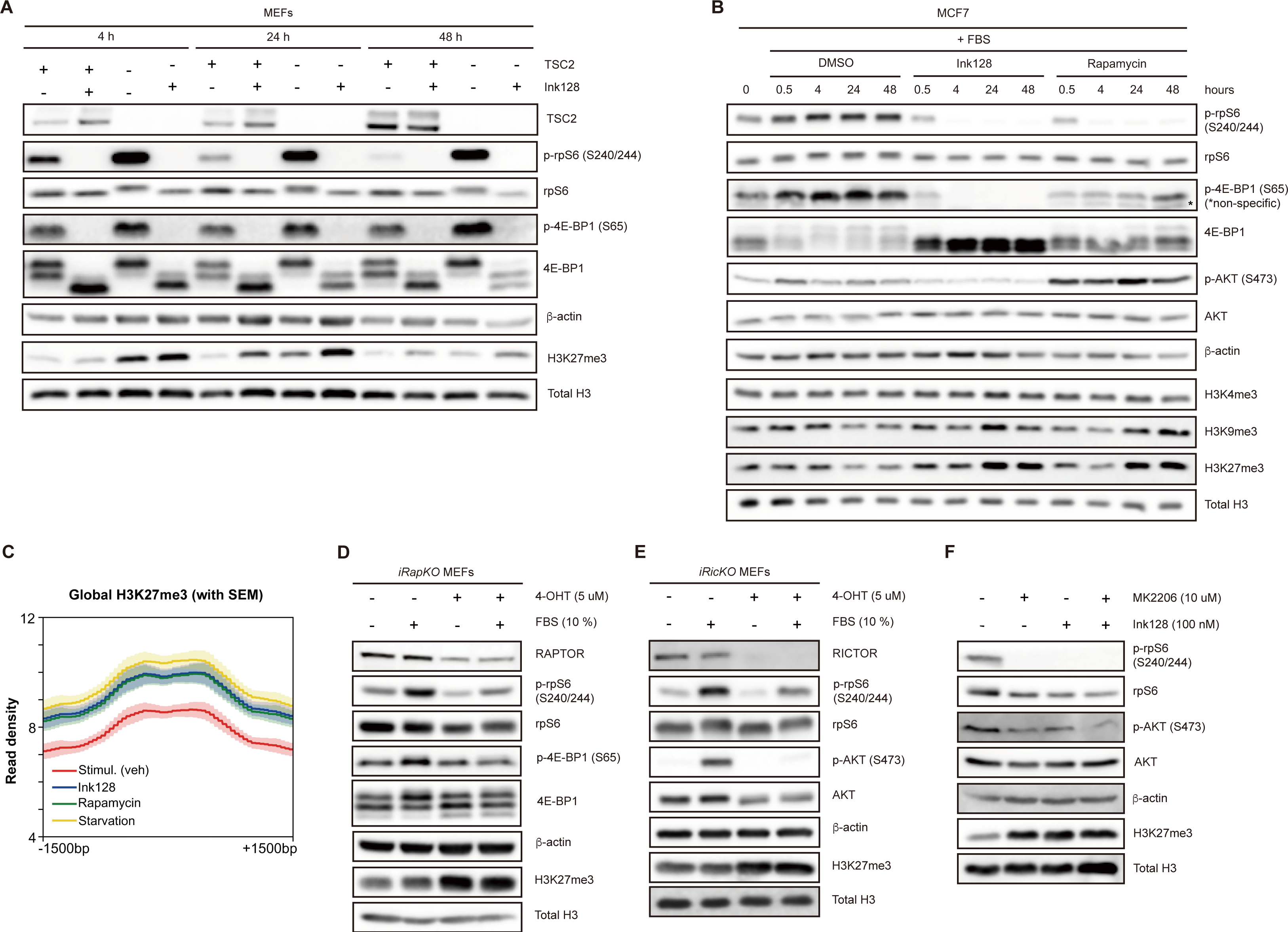
mTOR inhibition is accompanied by selective induction of H3K27me3 levels. (A) *TSC2* WT and KO MEFs were serum-starved overnight and then stimulated with 10% FBS in the presence of Ink128 (100 nM) for the indicated time periods. Levels of the indicated proteins were monitored by Western blotting (n=3). β-actin served as a loading control. (B) MCF7 cells were serum-starved overnight and then stimulated with 10% FBS in the presence of a vehicle (DMSO), Ink128 (100 nM) or rapamycin (50 nM) for the indicated periods. Levels of the indicated proteins were monitored by Western blotting (n=3). β-actin served as a loading control. (C) The normalized average read peak density profiles for H3K27me3 with SEM in MCF7 treated as indicated. (D-E) Levels of the indicated proteins in *iRapKO* (D) and *iRicKO* MEFs (E) were determined by Western blotting (n=3). Cells were serum-starved for 6 hours and then stimulated with 10% FBS for 30 min. β-actin was used as a loading control. (F) Levels of the indicated proteins in MCF7 cells treated as denoted for 48 hours were determined by Western blotting (n=3). β-actin was used as a loading control.

Considering the unexpected findings that the inhibition of mTOR by Ink128 did not cause a reduction, but rather induced H3K27me3 levels independently of the TSC2 status in cells (Figure 2A), we next sought to further characterize the effect of inhibition of mTOR activity on histone methylation across different cell lines. To this end, we utilized MCF7 (breast cancer cells) and HCT116 (colorectal cancer cells) harboring activating PI3K mutations, and consequently exhibiting elevated mTOR signaling. To modulate mTOR activity in these cells, we used rapamycin, which acts as an allosteric mTOR inhibitor that directly targets mTORC1 but not mTORC2, and Ink128 which targets active site of mTOR and suppresses both mTORC1 and mTORC2 ^22^. As above, MCF7 cells were serum starved overnight and then treated with Ink128, rapamycin or a vehicle (DMSO) in the presence of 10% FBS. These experiments showed that short-term mTOR inhibition (30 minutes and 4 hours) does not exert a major effect on H3K4me3, H3K9me3 or H3K27me3 levels (Figure 2B). However, when the Ink128 or rapamycin treatments were extended to 24 and 48 hours, increased levels of H3K9me3 and H3K27me3, but not H3K4me3, were observed relative to vehicle-treated cells, in which H3K9me3 and H3K27me3 levels were reduced at these time points (Figure 2B and Figure S1A). Comparable observations were made in HCT116 cells, wherein mTOR inhibitors induced H3K27me3 levels after 48h treatment as compared to vehicle treated controls (Figure S1B). Notably, in both MCF7 and HCT116 cells the effects of mTOR inhibitors on H3K27me3 were comparable to those observed when mTOR signaling was suppressed by serum starvation (Figure S1A-B). Using chromatin immunoprecipitation-sequencing (ChIP-seq), we confirmed that mTOR inhibition for 48 hours results in increased H3K27me3 levels in MCF7 cells (Figure 2C). Unlike H3K27me3, mTOR inhibition did not cause a noticeable difference in global H3K9me3 enrichment as monitored by ChIP-seq (Figure S2A). Furthermore, considering that H3K9me3 is typically situated at the nuclear periphery, albeit with a few focal points in the center of the nucleus ^34^, we employed immunofluorescence (IF) to quantify the number of H3K9me3 foci and measure the mean fluorescence intensity (MFI) (Figure S2B). We observed that Ink128 treatment did not affect the number of H3K9me3 foci relative to serum stimulated cells, whereas serum starvation and rapamycin treatment led to a slight decrease in the number of foci (Figure S2C). Moreover, the MFI of H3K9me3 normalized to DAPI intensity was lower in mTOR-inhibited cells compared to the control, serum stimulated cells (Figure S2D). These findings, together with the observation that mTORC1 activation selectively upregulates H3K27me3 levels, motivated us to focus on the effects of mTOR inhibition on H3K27me3.

### mTOR inhibition alters H3K27me3 deposition on a genome-wide scale

Next, we set out to establish the consequences of altered H3K27me3 levels upon mTOR inhibition on the steady-state mRNA levels. Combined analysis of ChIP-seq and RNA-seq data obtained from MCF7 cells after 48 hours of rapamycin or Ink128 treatment revealed a negative correlation between the two datasets which is consistent with H3K27me3 being associated with transcriptional repression (Figure S3A-C). We however found only a limited number of genes that exhibited congruent changes that were significant in both ChIP-seq and RNA-seq data sets (Figure S3D-E). We confirmed that the effects of mTOR inhibitors on H3K27me3 and *SOX2* gene expression are likely to be mediated by H3K27me3 using ChIP-qPCR and RT-qPCR (Figure S3F-H). Due to the limited number of genes that showed correlation between RNA-seq and ChIP-seq upon mTOR inhibition, we were unable to conduct further investigation into the pathway analysis of the set of differentially regulated genes. This limitation may arise from the fact that mTOR can also regulate other transcription factors, such as ATF4 and MYC ^26,35^. It is possible that the regulation of these transcriptional factors following mTOR inhibition also affects transcription, thereby making it challenging to identify genes that are specifically regulated via modulation of H3K27me3 levels. Additionally, it is possible that repetitive elements, which were filtered out of our RNA-seq analysis, may act as downstream effectors of mTOR signaling rather than coding genes.

### Inhibition of AKT, mTORC1, and mTORC2 induces H3K27me3 levels

To discern between the roles of mTORC1 and mTORC2 in regulating H3K27me3, we employed MEFs in which the mTORC1-specific component raptor (*iRapKO*) or the mTORC2-specific component rictor (*iRicKO*) can be depleted by 4-Hydroxytamoxifen (4-OHT) treatment ^36^. As compared to a vehicle control, 4-OHT treatment in *iRapKO* MEFs resulted in depletion of raptor levels (Figure 2D). This was accompanied by a decrease in mTORC1 activity as illustrated by a reduction in the phosphorylation of rpS6 (S240/244), and 4E-BP1 (S65) upon serum stimulation as compared to the control, serum stimulated cells that were not treated with 4-OHT (Figure 2D). Similarly, exposure of *iRicKO* MEFs to 4-OHT induced depletion of rictor and reduction of mTORC2 substrate AKT phosphorylation (S473) as compared to control, 4-OHT untreated and serum stimulated cells (Figure 2E). Of note, in these experiments, cells were stimulated with serum for only 30 minutes to test the effects of abrogation of raptor and rictor on mTORC1 and mTORC2 signaling, respectively. Importantly, 30-minute serum stimulation of MCF7 cells is not sufficient to affect H3K27me3 levels as compared to serum-starved cells (Figure 2B; DMSO treated cells). In turn, depletion of both raptor and rictor was accompanied by an increase in H3K27me3 levels as compared to control cells (Figure 2D-E). These findings suggest that the inhibition of both mTORC1 and mTORC2 may contribute to the induction of H3K27me3. In addition, treatment of MCF7 cells with the allosteric AKT inhibitor MK2206 resulted in downregulation of mTORC1 signaling as evidenced by the decrease in the phosphorylation of rpS6 (S240/244), which was paralleled by the H3K27me3 induction (Figure 2F). Therefore, prolonged inhibition of the AKT/mTOR axis is mirrored by the increase in H3K27me3 levels.

### mTOR inhibition-induced increase in H3K27me3 is largely independent of cell cycle

We next sought to dissect the mechanism(s) underpinning the role of mTOR inhibition in upregulation of H3K27me3 levels. *De novo* histone modifications are typically incorporated into chromatin during cell division ^37^, and mTOR inhibition results in cell cycle arrest ^38^. To this end, we explored the plausibility that observed increase in H3K27me3 may be secondary to the effects of mTOR inhibition on cell cycle. As expected, mTOR inhibition with either Ink128 or rapamycin slowed down G1/S progression (Figure S4A) and this coincided with an increase in H3K27me3 levels in MCF7 cells (Figure S4B). However, we did not observe a significant fluctuation in H3K27me3 levels across different cell cycle phases in either, serum starved, mTOR inhibitor or vehicle treated MCF7 cells (Figure S4C). This suggests that is unlikely that alterations in H3K27me3 levels are secondary to mTOR-dependent perturbations of cell cycle.

### mTOR inhibition-driven H3K27me3 is independent of metabolic perturbations

Emerging data indicate that perturbations in intermediate metabolites impact on histone methylation ^39^. For instance, α-KG serves as a key co-factor of histone and other demethylases ^40^. Therefore, we investigated whether metabolic perturbations caused by mTOR inhibition may explain the induction in H3K27me3 levels. Considering that succinate, fumarate, and 2-hydroxyglutarate (2-HG) are known to inhibit α-KG-dependent dioxygenases, including histone demethylases ^41,42^, we carried out gas chromatography–mass spectrometry (GC-MS) to measure the effects of mTOR inhibitors on the levels of these metabolites in MCF7 cells. Since Ink128 reduced the levels of succinate, fumarate, and 2-HG, and rapamycin reduced the levels of 2-HG compared to control, vehicle-treated cells (Figure 3A), we excluded the possibility that the observed increase in H3K27me3 levels was due to interference of these metabolites with α-KG-dependent demethylases. However, we also observed a significant decrease in α-KG levels upon mTOR inhibition (Figure 3A), which prompted us to hypothesize that mTOR inhibitors may be inducing H3K27me3 by reducing α-KG levels and consequently demethylase activity. To test this hypothesis, we supplemented MCF7 cells with 5 mM α-KG in the presence of mTOR inhibitors. This approach resulted in increase in intracellular α-KG (Figure 3B), but this was not paralleled by a conspicuous decrease in H3K27me3 levels in mTOR inhibitor-treated cells relative to vehicle treated controls (Figure 3C). In addition to α-KG, we also utilized dimethyl α-KG (DMKG), which is α-KG derivative with superior cell permeability relative to α-KG ^43^. Similar to the α-KG supplementation experiments, the addition of DMKG did not exhibit a major effect on induction of H3K27me3 that was triggered by mTOR inhibition (Figure 3D). Overall, our results imply that the hypermethylation of H3K27me3 following mTOR inhibition is independent of the alterations in α-KG levels, or levels of metabolites that interfere with demethylase activity.

**Figure 3.**
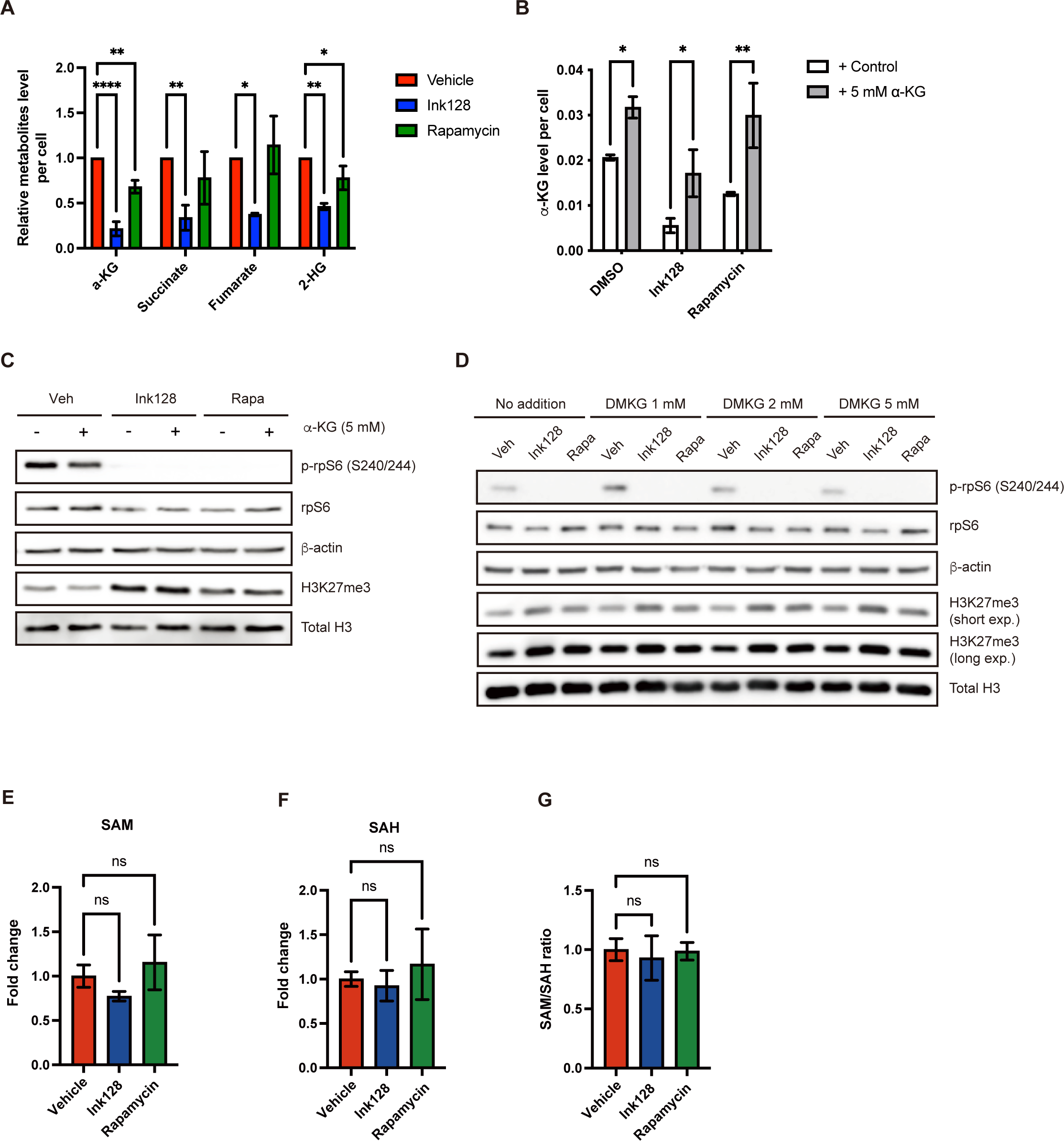
mTOR inhibition-driven H3K27me3 is not secondary to metabolic perturbations. (A) Quantification of steady-state levels of metabolites from MCF7 cells that were serum-starved overnight and then stimulated with 10% FBS in the presence of a vehicle (DMSO), Ink128 (100 nM) or rapamycin (50 nM) for 48 hours. Metabolites were extracted, profiled by GC-MS, and normalized to cell numbers. Bars represent mean ± SD (n=3). ns: no significance, *P< 0.05, **P<0.01, ****P< 0.0001, one-way ANOVA with Dunnett’s post-test compared to 10% FBS stimulated, vehicle (DMSO) treated cells. (B) Quantification of steady-state levels of α-KG from MCF7 cells treated with 5 mM α-KG in the presence of 100 nM Ink128, 50 nM rapamycin or vehicle (DMSO) for 48 hours. Metabolites were extracted, profiled by GC-MS, and normalized to cell numbers. Bars represent mean ± SD (n=2). *P< 0.05, **P<0.01, paired t-test compared to the control without addition of α-KG. (C-D) Levels of the indicated proteins in MCF7 treated with 5 mM α-KG (C) or indicated concentration of DMKG (D) in the presence of 100 nM Ink128, 50 nM rapamycin or vehicle (DMSO) for 48 hours were determined by Western blotting (n=3). β-actin served as a loading control. (E-F) Quantification of SAM and SAH levels at steady state from MCF7 cells that were serum-starved overnight and then stimulated with 10% FBS in the presence of a vehicle (DMSO), Ink128 (100 nM), or rapamycin (50 nM) for 48 hours. Metabolites were extracted, profiled by LC-MS, and normalized to cell numbers. ns: no significancy. Bars represent mean ± SD (n=3) (G) The SAM/SAH ratio was calculated from Figure 5E-F. Bars represent mean ± SD (n=3)

We next examined the effects of mTOR inhibitors on levels of SAM and S-Adenosyl-L-homocysteine (SAH) in MCF7 cells. SAM and SAH are crucial substrates for methyltransferases that are vital for their enzymatic function ^44^. These experiments revealed that mTOR inhibition had no significant impact on the levels of SAM or SAH in MCF7 cells (Figure 3E-F), and the SAM/SAH ratio remained relatively unaffected by mTOR inhibitors (Figure 3G). These findings suggest that the availability of methyl donors is unlikely to be the key factor in mediating the effects of mTOR inhibitors on H3K27me3. Collectively, these findings suggest that metabolic perturbations affecting α-KG, SAM or SAH levels are not sufficient to explain the induction of H3K27me3 levels upon mTOR inhibition.

### mTOR inhibition-driven H3K27me3 is unlikely to be mediated by KDM6A and KDM6B demethylases

Next, we investigated the implication of key regulators of H3K27 methylation in mediating the effects of mTOR inhibition on this histone modification. KDM6A and KDM6B are recognized as primary demethylases for H3K27me3 ^45^. In turn, EZH1 and EZH2 function as the catalytic subunit of PRC2, responsible for depositing H3K27me3 ^46^. Firstly, we studied the impact of mTOR inhibition on the RNA and protein levels of H3K27 demethylases, KDM6A and KDM6B. In these experiments, MCF7 cells were serum starved overnight and then stimulated with 10% FBS in the presence of the vehicle, rapamycin or Ink128 or kept starved for 48 hours. Serum starvation and Ink128 treatment significantly induced *KDM6A* and *KDM6B* mRNA levels relative to serum stimulated, vehicle treated cells (Figure 4A). KDM6A protein levels were however only modestly increased in response to Ink128 treatment, but not serum starvation as compared to serum stimulated cells (Figure 4B). Rapamycin treatment failed to significantly affect RNA and protein levels of H3K27 demethylases (Figure 4A-B). Lastly, KDM6B protein levels remained largely unchanged across all conditions (Figure 4B). These findings suggest that the increase in H3K27me3 observed upon mTOR inhibition cannot be explained by a reduction in KDM6A and/or KDM6B levels. We then sought to investigate whether mTOR inhibition may affect the activity of KDM6A and KDM6B by inducing their subcellular re-distribution. However, mTOR inhibition in MCF7 cells did not result in appreciable effects on the localization of either KDM6A or KDM6B (Figure 4C). Altogether, these findings suggest that KDM6A and KDM6B are not likely to mediate the effects of mTOR inhibitors on H3K27me3.

**Figure 4.**
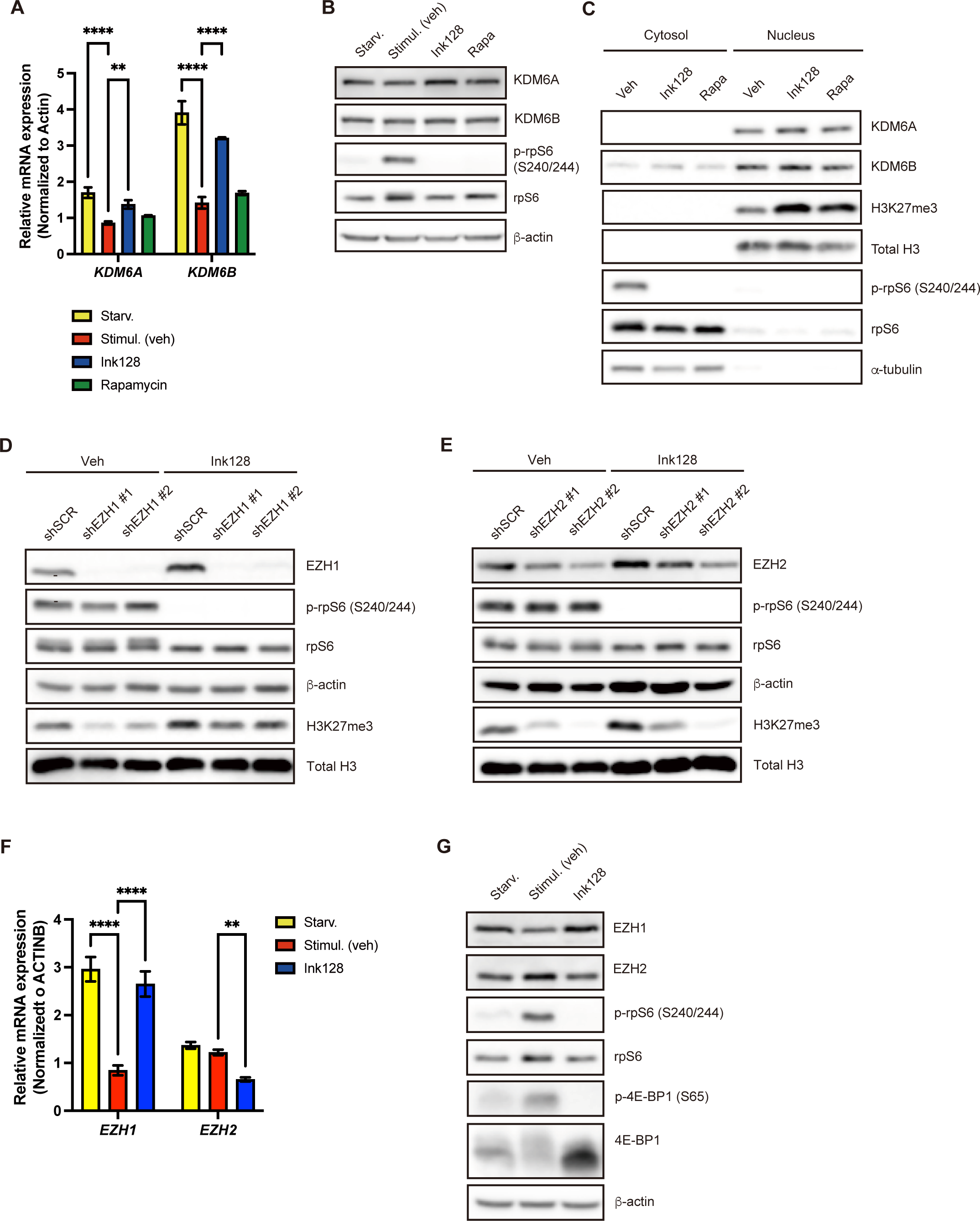
The increase in H3K27me3 upon mTOR inhibition correlates with EZH1 upregulation and is independent of KDM6A and KDM6B demethylases. (A) The levels of *KDM6A* and *KDM6B* mRNAs in MCF7 cells that were serum-starved overnight and then stimulated with 10% FBS in the presence of a vehicle (DMSO Stimul (veh)-red), Ink128 (100 nM; Ink128-blue) or rapamycin (50 nM; rapamycin-green), or were kept serum-starved (Starv-yellow) for 48 hours were determined by RT-qPCR. Bars represent mean ± SD (n=3). **P<0.01, ****P< 0.0001, one-way ANOVA with Dunnett’s post-test (B) Levels of the indicated proteins in MCF7 upon the treatment with DMSO, 100 nM Ink128, or 50 nM rapamycin for 48 hours were determined by Western blotting (n=3). β-actin served as a loading control. (C) Immunoblot analysis of cytoplasmic and nuclear extracts from MCF7 cells treated with 100 nM Ink128 or 50 nM rapamycin for 48 hours (n=3). α-tubulin and H3 served as loading controls of cytoplasmic and nuclear proteins, respectively. (D) Levels of the indicated proteins in HCT116 cells infected with a scrambled (shSCR) or EZH1-specific shRNA (shEZH1) and treated with a vehicle (DMSO) or Ink128 (100 nM) were determined by Western blotting (n=3). β-actin served as a loading control. (E) Levels of the indicated proteins in HCT116 infected with a scrambled (shSCR) or EZH2-specific shRNA (shEZH2) treated with a vehicle (DMSO) or Ink128 (100 nM) were determined by Western blotting (n=3). β-actin served as a loading control. (F) The levels of *EZH1* and *EZH2* mRNAs in MCF7 cells that were serum-starved overnight and then stimulated with 10 % FBS in the presence of a vehicle [DMSO; Stimul (Veh)-red] or Ink128 (100nM; Ink128-blue), or were kept serum-starved (Starv-yellow) for 48 hours were determined by RT-qPCR. **P<0.01, ****P< 0.0001, one-way ANOVA with Dunnett’s post-test (n=3) (G) Levels of the indicated proteins in MCF7 that were serum-starved overnight and then treated with a vehicle (DMSO) or Ink128 (100 nM) in the presence of 10% FBS, or were kept serum-starved for 48 hours were determined by Western blotting (n=3). β-actin served as a loading control.

### EZH1/2 ratio is altered when mTOR is inhibited

We next investigated whether PRC2 is accountable for the induction of H3K27me3 upon mTOR inhibition. We first depleted EZH1 or EZH2 in HCT116 cells using shRNA. These cells were then treated with Ink128 for 48 hours. As expected, the levels of H3K27me3 decreased in cells depleted of EZH1 or EZH2 as compared to control, scrambled shRNA infected cells (Figure 4D-E). The increase in H3K27me3 induced by Ink128 treatment was attenuated by depletion of either EZH1 or EZH2 (Figure 4D-E), highlighting the cooperativity between these epigenetic writers. Next, we sought to dissect the potential mechanisms that may govern EZH1 and EZH2 activity in the context of mTOR inhibition. We first evaluated the effects of suppression of mTOR signaling on EZH1 and EZH2 mRNA and protein levels by serum starving MCF7 cells for 48 hours and then stimulating them with 10% FBS in the presence of a vehicle or Ink128. These experiments revealed that relative to 10% FBS stimulation, serum starvation and Ink128 increased levels of *EZH1* mRNA and protein while decreasing the levels of EZH2 protein (Figure 4F-G). This increase in EZH1 levels in serum-starved and Ink128 treated cells may serve as a compensatory response to the decrease in EZH2 abundance to maintain activity of PRC2 ^47^. Notably, the treatments did not strongly impact *EZH2* mRNA levels, which were modestly downregulated only in Ink128, but not in serum starved cells (Figure 4F). This aligns with previous result showing that mTORC1 regulates EZH2 levels via 4E-BPs-dependent translational mechanisms ^23^. Altogether, these results suggest that the increase in H3K27me3 in response to mTOR inhibition may be mediated by EZH1-dependent effects on PRC2 activity. This is opposite to constitutive mTORC1 activation wherein the increase in H3K27me3 appears to be chiefly mediated by EZH2.

We also investigated whether mTOR may affect H3K27me3 by altering the assembly of PRC2. Ink128 however did not affect the association of EZH2 with other PRC2 components in MCF7 cells, which suggests that the effects of mTOR inhibition-induced increase in H3K27me3 are not mediated via the effects on PRC2 assembly (Figure S5A). Finally, H3K9 methyltransferases GLP and G9a have been associated with PRC2-mediated modifications of H3K27me2 and H3K27me3. GLP plays a role in the formation of H3K27me2, while the enzymatic activity of G9a influences the genomic recruitment of PRC2 to a specific group of its target genes ^48,49^. Thus, we exploited the plausibility that mTOR inhibitors may induce H3K27me3 indirectly via GLP and/or G9a. However, mTOR inhibition by Ink128 did not affect G9a, while decreasing GLP protein levels relative to the control, vehicle treated MCF7 cells (Figure S5B). Furthermore, GLP/G9a inhibition with UNC0642 failed to attenuate the induction of H3K27me2 and H3K27me3 caused by Ink128 (Figure S5C). These findings suggest that the changes in H3K27me3 induced by mTOR inhibition are unlikely to involve GLP/G9a. Altogether, our data show that the induction of H3K27me3 upon mTOR inhibition alters the EZH1/2 ratio that may be required to maintain PRC2 activity under mTOR inhibition. In turn, the effects of mTOR inhibition on H3K27me3 levels do not appear to be mediated by modulation of PRC2 assembly or GLP/G9a.

### The anti-proliferative effects of mTOR inhibitors are partially mediated via H3K27me3

We next sought to establish the role of H3K27me3 in mediating the effects of mTOR inhibitors on cell functions. Herein, we focused on well-established anti-proliferative effects of mTOR inhibitors ^38^. Indeed, in line with previous findings ^38^, we observed a decrease in cell proliferation upon mTOR inhibition in both MCF7 and HCT116 cells (Figure S6A-B). Suppression of PRC2 activity by GSK126 also reduced proliferation of MCF7 cells (Figure S6C). We then explored the effects of mTOR inhibitors on cell proliferation under conditions wherein H3K27me3 levels are reduced by GSK126. Strikingly, although each inhibitor alone exerted anti-proliferative effects, GSK126 attenuated the anti-proliferative effects of Ink128 in MCF7 cells (Figure 5A). These findings indicate that inhibition of H3K27me3 hinders the anti-proliferative effects of mTOR inhibitors. To further investigate this, we used the DIPG13 cell line, which carries a heterozygous *H3K27M* mutation ^50^, along with DIPG13 *H3K27M*-KO cells, wherein the mutant allele was removed using CRISPR/Cas9 technology ^51^. Consistent with the dominant-negative effect of the *H3K27M* mutation on H3K27me3 levels ^52^, we failed to detect appreciable levels of H3K27me3 in *H3K27M* DIPG13 cells, whereby H3K27me3 was rescued in DIPG13 *H3K27M*-KO cells (Figure 5B). Ink128 failed to induce H3K27me3 in *H3K27M* DIPG13 cells, while increasing H3K27me3 in DIPG13 *H3K27M-*KO cells (Figure 5B). Consistent with the experiments wherein PRC2 function was disrupted pharmacologically, the anti-proliferative effects of Ink128 were attenuated in DIPG13 *H3K27M* relative to DIPG13 *H3K27M-*KO cells (IC50 32.59 vs. 9.35; Figure 5C). In conclusion, these findings suggest that the anti-proliferative effects of mTOR inhibitors are at least in part mediated by H3K27me3.

**Figure 5.**
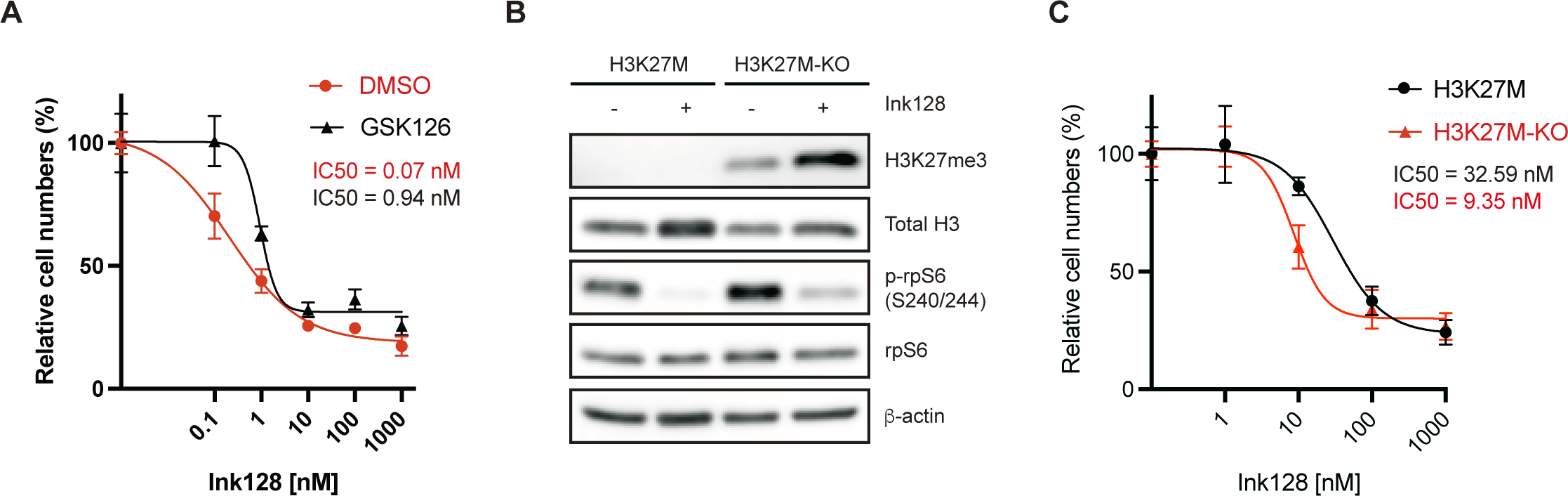
The effect of mTOR inhibition on cell proliferation is partially mediated by induction of H3K27me3. (A) MCF7 cells were treated with indicated concentrations of Ink128 in the presence of DMSO (vehicle, red) or 2 uM GSK126 (black) for 72 hours. Data are presented as means ± SD at each time point (n=3). The numbers of viable cells at each time point were determined using the cell counter, and IC_50_ was calculated by GraphPad. (B) Levels of the indicated proteins in DIPG13 *H3K27M* and *H3K27M*-KO cells were determined by Western blotting (n=2). DIGP13 cells were treated with 100 nM Ink128 or vehicle (DMSO) for 48 hours. β-actin served as a loading control. (C) DIPG13 *H3K27M* (black) and *H3K27M*-KO cells (red) were treated with indicated concentrations of Ink128 for 72 hours. Data are presented as means ± SD at each time point (n=3). The numbers of viable cells at each time point were determined using the cell counter, and IC_50_ was calculated by GraphPad.

## Discussion

In this study, we explored the impact of mTOR on histone methylation. We observed that paradoxically, both mTOR activation and inhibition induce H3K27me3, a gene-repressive mark. In line with previous reports ^53^, we observed no changes in H3K4me3 levels upon mTOR inhibition. This suggests that the effects of mTOR on histone methylation are selective, whereby H3K27me3 is induced while other histone methylation marks (e.g., H3K4me3 or H3K9me3) are not considerably affected. Opposite to this report, previous studies ^29,54^ failed to detect H3K27me3 induction upon mTOR inhibition. These discrepancies could stem from the duration of mTOR inhibition and the diversity of experimental models, including different cell types. Considering the relatively slow turnover of histone modification ^55^, it is likely that the effects of modulation of mTOR on H3K27me3 may not be apparent in acute treatments. Indeed, our time course studies revealed that long-term treatments (∼24-48 hours) with mTOR inhibitors were required to observe alteration in H3K27me3 levels in MEFs, MCF7 (Figure 2A-B) and HCT116 (Figure S1B). In turn, prolonged mTOR inhibitor treatments on histone methylation may be secondary to the effects of mTOR modulation on cell functions. We, however, provide evidence that the observed effects of mTOR inhibitors on H3K27me3 are unlikely to be secondary to cell cycle or metabolic perturbations.

While we confirmed that mTORC1 hyperactivation in various models induces EZH2 protein level in a 4E-BPs-dependent manner ^23^, mechanistic underpinnings governing the increases in H3K27me3 driven by mTOR inhibition did not appear to involve an increase in EZH2 levels. Indeed, despite induction of H3K27me3 upon mTOR inhibition, both serum starvation and Ink128 decreased EZH2 levels, which is consistent with the activation of 4E-BPs and suppression of EZH2 protein synthesis ^23^. In turn, these treatments were associated with the induction of EZH1 mRNA and protein levels, thus suggesting that in the context of mTOR inhibition, EZH1 may be compensating for a decrease in EZH2 to maintain PRC2 activity and sustain deposition of H3K27me3. Nonetheless, the precise mechanisms of the induction of H3K27me3 caused by mTOR inhibition remains uncertain. Several mechanisms that were not tested in the present study may be triggered by mTOR inhibition. Firstly, mTOR may engage other PRC2 components in addition to EZH1 and EZH2. Although SUZ12 and EED could not explain the increase in H3K27me3, as both are reduced by Ink128 treatment (Figure S5A), additional PRC2 components including RBBP4, and other accessory factors (JARID2, AEBP2, EPOP and PCLs), that are generally thought to modulate PRC2 function may be involved ^56^. Moreover, the increased levels of H3K27me3 resulting from mTOR inhibition could potentially involve the modulation of other histone modifications that enhance H3K27me3, such as PRC1-mediated H2AK119ub1 ^57^, nuclear S6K1-induced H2BS36 phosphorylation ^58^, and NSD1-mediated H3K36 methylation ^59–61^. Lastly, given the reported role of mTOR in the nucleus ^62,63^, it is conceivable that nuclear mTOR may interact with H3K27me3 modifiers and/or histones, prompting H3K27me3 induction during mTOR inhibition. Altogether, we believe that our initial findings suggest that future studies are warranted to establish the precise mechanisms of the role of mTOR inhibition in the regulation of H3K27me3.

Importantly, reducing H3K27me3 levels attenuated the anti-proliferative effects of mTOR inhibition, thus suggesting that this histone modification may represent a previously unrecognized mediator of the anti-proliferative effects of mTOR inhibitors. Intriguingly, H3K27me3 levels were also induced upon constitutive activation of mTORC1 in *TSC2* KO MEFs, whereby mTOR hyperactivation is generally linked to a pro-proliferative cellular state ^38^. Although this remains to be established, the plausible resolution of these apparently contradictory findings may stem from the differences in the mechanisms of induction of H3K27me3 levels under conditions of mTOR activation vs. inhibition. To this end, it is likely that the cohorts of genes affected by changes in H3K27me3 may differ significantly between the cells in which mTOR is activated vs, those in which it is suppressed.

mTOR activity is frequently upregulated across a broad range of malignancies ^64^. H3K27me3 dysregulation is also common in cancer, with aberrant EZH2 levels observed in a variety of neoplasia ^65–70^. Indeed, both epigenetic alterations and mTOR-dependent translational reprograming have been shown to induce metabolic and phenotypic plasticity of cancer cells ^71–73^. This suggests that the cross-talk between mTOR and epigenetic mechanisms may play a role in cancer cell plasticity which has been implicated in tumor progression and poor therapeutic responses ^74^. Accordingly, both mTOR and PRC2 inhibitors exhibit anti-neoplastic properties, whereby these compounds are either in clinical trials or already employed in the clinic in oncological indications ^75–78^. Although our studies were limited to cell culture, they suggest that the anti-proliferative activity of mTOR inhibitors may be attenuated under conditions of restrained H3K27me3. This may have significant implications for therapeutic strategies combining mTOR inhibitors and H3K27me3 modulators, and application of mTOR inhibitors in tumors carrying *H3K27M* mutations.

In summary, this study provides evidence suggesting that mTOR may selectively influence histone modifications. Both constitutive mTOR activation and mTOR inhibition resulted in increased H3K27me3 levels, while H3K4me3 and H3K9me3 abundance remained largely unaffected. Furthermore, we show that induction of H3K27me3 by mTOR inhibitors is required for their anti-proliferative effects. Finally, these results emphasize the importance of studying the cross-talk between mTOR and epigenetic mechanisms in the context of different cell functions.

### Limitations of the study

Although we undertook a number of approaches to dissect molecular underpinnings of H3K27me3 induction upon mTOR inhibition, most of these efforts resulted in negative data. To this end, the major limitation of our study is the lack of precise mechanism(s) linking mTOR inhibition to H3K27me3 induction. Moreover, our experiments were limited to cell culture, and thus it is important to establish the robustness of reported observations in *in vivo* models. Our study was also limited to employing *TSC2*-null MEFs to model constitutively activated mTORC1. Therefore, it is necessary to confirm our findings in other models of mTORC1 hyperactivation including overexpressing constitutively active RagA and/or RagC ^79,80^.

## Data availability

ChIP/RNA-seq data is available in NCBI’s Gene Expression Omnibus (GEO; http://www.ncbi.nlm.nih.gov/geo) accession number GSE246889. This article does not report original code. Any additional information required to reanalyze the data reported in this article is available from the lead contact on request.

## Acknowledgments

We are thankful to Shannon McLaughlan and Dr. Laura Lee for technical assistance and to the members of Postovit’s, Witcher’s, and Topisirovic’s laboratories for helpful discussions. We would like to thank Dr. Nada Jabado for providing DIPG13 cells, carrying the *H3K27M* mutation, and a paired set of DIPG13 *H3K27M*-KO cells. We would also like to express our appreciation to Dr. David Kwiatkowski and Dr. Michael N. Hall for providing *TSC2* WT/KO MEFs and *iRapKO/iRicKO* MEFs, respectively. This research was funded by the Terry Fox Foundation (TFF) Oncometabolism Team Grant TFF-242122 to IT and Canadian Institutes for Health Research (CIHR) PJT-183843 to IT. HK and PJ are supported by Fonds de Recherche du Québec – Santé (FRQS) Fellowships. DP is supported by CIHR Postdoctoral Fellowship and Cancer Research Society (CRS) The Next Generation of Scientists Award (NGS). IT is supported by FRQS Senior Investigator award. MW is supported by the CIHR, CRS and holds a salary award from FRQS. JUS is supported by CIHR PJT-103526. LMP is supported by the Canadian Cancer Society (CCS) Innovation Grants (CCSRI). Metabolic analysis was performed at The Rosalind and Morris Goodman Cancer Research Centre’s Metabolomics Core Facility, which is supported by the Canada Foundation for Innovation, The Dr. John R. and Clara M. Fraser Memorial Trust, the Terry Fox Foundation (TFF Oncometabolism Team Grant; TFF-242122) and McGill University.

## Author contributions

HK conceived the study and performed experiments; BL and FA helped with ChIP/RNA-seq experiments and analyzed ChIP/RNA-seq data; DP, DA and ML helped performing metabolomic studies, LMP, MW and IT participated in conceiving and conceptualizing the study and provided supervision and funding. PJ, JUS, and MM provided key reagents. All authors participated in writing the paper.

## Inclusion and diversity

We endorse conducting research in an inclusive, diverse, and equitable manner.

## STAR★METHODS

### RESOURCE AVAILABILITY

#### Lead contact

Further information and requests for reagents may be directed to, and will be fulfilled upon reasonable request by, the lead contact Ivan Topisirovic.

#### Materials availability

We generated a breast cancer cell line isolated from MMTV-PyMT/*Eif4ebp1/2*-null mice which is available upon request.

#### Data and code availability

- RNA-seq and ChIP-seq data have been deposited at GEO and are publicly available as of the date of publication. Accession numbers are listed in the key resources table. Original Western blot images have been deposited at Mendeley and are publicly available as of the date of publication. The DOI is listed in the key resources table.
- This paper does not report original code.
- Any additional information required to reanalyze the data reported in this paper is available from the lead contact upon reasonable request.

### EXPERIMENTAL MODEL AND SUBJECT DETAILS

#### Cell cultures

*TSC2* WT and KO MEFs were obtained from Dr. Kwiatkowski’s lab ^31^. The breast cancer cell lines were derived from MMTV-PyMT (mouse mammary tumour virus-Polyoma Virus middle T antigen)/*Eif4ebp1/2*-null mice. *MMTV-PyMT* mice ^81^ were crossed with *Eif4ebp1-/-/Eif4ebp2-/-mice* ^82^ in the FVB background. A tumor cell line was established after explanting the breast tumor tissue of a female mouse at necropsy into cell culture. Subsequent re-expression of 4E-BP1 form was performed through retroviral transduction of the pBABE-puro construct. These cells were cultured in DMEM medium supplemented with 2.5% heat-inactivated FBS, 100 UI/ml penicillin/streptomycin, 50 ug/ml gentamicin and 1% mammary epithelial growth supplement made in house. MCF7 and HCT116 were obtained from ATCC. MCF7 and HCT116 were cultured in RPMI-1640 and DMEM respectively, supplemented with 10% heat-inactivated FBS, 1% penicillin/streptomycin, and 2 mM l-glutamine. Inducible raptor (*iRapKO*) and rictor (*iRicKO*) knockout MEFs were provided by Dr. Hall’s lab ^36^. Cells were treated with 5 uM of 4-hydroxytamoxifen (4-OHT) for at least three days. The tumor-derived cell line DIPG13, carrying the *H3K27M* mutation, and a paired set of DIPG13 *H3K27M*-KO cells were provided by Dr. Jabado’s lab ^83^. Cells maintained in Neurocult NS-A proliferation media supplemented with bFGF (10 ng/ml), rhEGF (20 ng/ml), and heparin (0.0002%) on plates coated in poly-L-ornithine (0.01%) and laminin (0.01 mg/ml). For all cell lines other than DIPG13, 0.05% Trypsin-EDTA was utilized to dislodge the cells from the plate. For DIPG13 cells, accutase was used to detach cells from the plate. Cells were grown in a humidified environment at 37 °C with 5% CO_2_. All the reagents are listed in Key resources table.

## METHOD DETAILS

### Lentiviral packaging and infection

The transfection of lentiviral constructs was performed using jetPRIME transfection agent according to the manufacturer’s protocol (Polyplus transfection). HEK293T cells were co-transfected with 3 µg of target shRNA-containing plasmid (human shEZH1, human shEZH2, mouse shEzh2, and shScramble), 2 µg of psPAX2 packaging plasmid, and 1 µg of pMD2.G plasmid. The media was changed 24 hours later and collected 48 hours post-transfection. The virus-containing media were filtered through a 0.45 µm filter (ThermoFisher Scientific) and mixed with fresh media at a 1:1 ratio before adding to pre-seeded target cells. To enhance transduction efficiency, 4 µg/ml of polybrene (MilliporeSigma) was added to the cells. The cells were infected in two rounds, with a single infection per day. After 24 hours of last infection, selection was performed using 2 µg/ml of puromycin (Bio Basic) to collect cells expressing the desired shRNAs. The list of reagents and shRNAs are described in Key resources table.

### Western blotting

Cells were washed with ice-cold PBS twice and lysed with RIPA buffer (1% NP40, 0.1% SDS, 50 mM Tris-HCl pH 7.5, 150 mM NaCl, 0.5% sodium deoxycholate, 1 mM PMSF, 1 mM DTT, 1X PhosSTOP and 1X proteinase inhibitor cocktail). The lysates were sonicated with a probe sonicator (Fisher Scientific Sonic Dismembrator Model 500) for 4 sec × 2 times at 30% power and clarified at 4 °C (10 min at 16,000 g). Protein concentrations in the supernatants were determined using BCA™ kit (ThermoFisher Scientific). Samples were boiled in 5× Laemmli buffer at 95 °C for 5 min, proteins were separated by SDS-PAGE and transferred using wet mini-transfer system omniBLOT Complete Systems (Cleaver scientific) onto nitrocellulose membranes. The membranes were blocked in 3% skim milk w/v in TBST buffer (0.1% Tween 20 in 1× TBS) and then incubated with primary antibodies, which were prepared in 3% BSA in TBST at 4 °C. Membranes were washed with TBST (3 × 10 min) and incubated for 1 hour with HRP-conjugated secondary antibodies, which were prepared in 3% skim milk/TBST. After washing the membranes with TBST (3 × 15 min), specific protein bands were revealed by chemiluminescence using ECL™ (BioRad) reagent on the Azure 600 (Azure Biosystems). When possible, the membranes were stripped and re-probed by the next set of the antibodies. This, however, was not possible in all cases because of the high protein abundance and/or strong intensity of some of the antibodies used. In the latter case the samples were run in parallel on separate gels. All the western blots were carried out in at least three independent replicates. The list of antibodies and dilutions that were used are described in Key resources table.

### Cell proliferation assay

For cell proliferation curves, cells were seeded in 6-well plates and incubated overnight. The media were replaced with treatment media containing 100 nM Ink128, 50 nM rapamycin or DMSO as a vehicle control. Every 24 hours, treatment media were aspirated, and the cells were trypsinized. Complete media were added to stop the trypsinization. Samples were collected, stained with trypan blue to exclude dead cells, and counted using an automated cell counter (Invitrogen). Data was analyzed using GraphPad Prism 9 software.

For the IC_50_ curves, cells were seeded in technical duplicates in 6-well plates and incubated overnight. The media were replaced with treatment media containing Ink128 at increasing concentrations for 72 hours. DMSO served as a control for the treatment. After 72 hours, cells were collected and counted using a countess automated cell counter (Invitrogen). IC_50_ curves were plotted, and the numerical values were computed using GraphPad Prism 9 software. All the cell counting assays were carried out in at least three independent replicates, each consisting of 2 technical replicates.

### LC-MS

Steady state SAM and SAH levels were measured by employing LC-MS/MS at the Metabolomics Core Facility of the Goodman Cancer Research Centre. After treatment with mTOR inhibitors (Ink128 100 nM or rapamycin 50 nM) or a vehicle (DMSO) for 48 hours, cells were washed in ammonium formate three times, then quenched in cold 50% methanol (v/v) and acetonitrile. Cells were lysed following bead beating at 30 Hz for 2 min. Cellular extracts were partitioned into aqueous and organic layers following dichloromethane treatment and centrifugation. Aqueous supernatants were dried down using a refrigerated speed-vac. Dried samples were subsequently resuspended in 50 μl of water. 5 μl of sample was injected onto an Agilent 6470 Triple Quadrupole (QQQ)-LC-MS/MS for targeted metabolite analysis of SAM and SAH. The liquid chromatography was performed using a 1290 Infinity ultra-performance binary LC system (Agilent Technologies, Santa Clara, CA, USA), with the following parameters: column and autosampler temperatures were 10 °C and 4 °C, respectively; flow rate of 0.6 ml/min with a Intrada Amino Acid column 3 μm, 3.0×150mm (Imtakt Corp, JAPAN). The gradient started at 100% mobile phase B (0.3% formic acid in acetonitrile) with a 3 min gradient to 27% A (100 mM ammonium acetate in 80% water / 20% ACN), followed by a 19.5 min gradient to 100% A. This was followed by a 5.5 min hold time at 100% mobile phase A, a 1 min gradient to 100% mobile phase B, and a subsequent re-equilibration time (7 min) before subsequent injection. The mass spectrometer was equipped with an electrospray ionization (ESI) source, and the samples were analyzed in positive mode. For each quantitated metabolite, multiple reaction monitoring (MRM) transitions were optimized using standards. Transitions for quantifier and qualifier ions were as follows: SAM (399.1 → 136.1 and 399.1 → 97) and SAH (385.1 → 136 and 385.1 → 250.1). Relative concentrations were determined by comparing the sample area under the curve to external calibration curves prepared in water. No adjustments were made for ion suppression or enhancement. The data were analyzed using MassHunter Quant (Agilent Technologies).

### GC-MS

After 48h treatment with mTOR inhibitors (Ink128 100 nM or rapamycin 50 nM) or a vehicle, the plates were quickly placed on ice. Cells were washed three times with chilled isotonic saline solution. Subsequently, 300 µL of 80% methanol pre-chilled to −20 °C was added to the cells. Cells were scraped from the wells and transferred to microcentrifuge tubes pre-chilled to −20 °C. 300 µL more of the 80% methanol was added to the leftover cells in the wells, and then scraped, collected, and pooled with the previously collected 300 µL fraction. The cell suspensions were lysed using a Diagenode Bioruptor sonicator (Diagenode Inc.) at 4 °C. The sonication was performed for 10 min with a 30-sec on-off cycle, using the high-power setting. The process was repeated three times to ensure complete recovery of metabolites. After centrifugation (16,000 g, 4 °C), the cell debris was discarded, and the supernatants were transferred to pre-chilled tubes. Subsequently, the supernatants were dried overnight at 4 °C in a CentriVap cold trap (Labconco) to remove the solvent. Dried pellets were dissolved in 30 µL of pyridine containing methoxyamine-HCl (10 mg·ml^−1^) (MilliporeSigma) using a sonicator and vortex. Samples were incubated for 30 min at 70 °C and then transferred to GC-MS injection vials containing 70 µL of N-tert-butyldimethylsilyl-N-methyltrifluoroacet-amide (MTBSTFA). Sample mixtures were further incubated at 70 °C for 1 hour. For GC–MS analysis, a volume of one microlitre was injected for each sample. GC–MS methods were conducted as previously described ^84^. Data analyses were executed using Agilent ChemStation and MassHunter software (Agilent). Each metabolite was normalized to the peak intensity of myristic acid-D27, and cell numbers were derived from cells seeded in parallel and identical conditions to those collected for GC-MS steady-state analysis. Data were expressed as fold change relative to vehicle (DMSO)-treated cells.

### Isolation of RNA and RT-PCR analysis

Total RNA was extracted from cells using Trizol (Ambion) following the manufacturer’s instructions. cDNA was made from purified total RNA using SensiFAST™ cDNA Synthesis Kit (#BIO-65053, Bioline) as per manufacturer’s protocol. The qPCR was conducted using SensiFAST™ SYBR® Lo-ROX kit (Bioline) on the AB7300 machine and analyzed using the 7300 system sds Software (Applied Biosystems). Activation 95 °C 10 min; Melting 95 °C 15 s; Annealing/extension 55 °C 1 min. Return to step 2 for 40 total cycles. Melting curve analysis was performed to ensure the production of a single amplicon. mRNA expression was measured by using relative standard method according to ABI Prism User Bulleting # 4. Each experiment was performed in independent triplicate, each consisting of a technical triplicate. Primers were designed using Primer3 (https://primer3.ut.ee/) for human genes. Primers for reactions are outlined in Key resources table.

### RNA sequencing and processing

Total RNA was extracted according to the Sigma RNA Extraction Kit (#RTN350-1KT, Sigma) protocol. RNA was sent to SickKids Genome centre for poly(A) RNA library preparation, using the NEBNext Ultra II Directional RNA Library Prep Kit for Illumina and sequencing of 50 M 100-bp paired-end reads per replicate on the Illumina NovaSEq 6000 platform. The quality of reads and sequencing was assessed both before and after trimming using the FastQC package (Babraham Bioinformatics). Before mapping, the reads were subjected to trimming using Trimmomatics ^85^ under the following conditions: ILLUMINACLIP:$Adapters:2:30:10:8:true, HEADCROP:4, SLIDINGWINDOW:4:30, LEADING:3, TRAILING:3, MINLEN:30. Subsequently, alignment to the hg19 human genome was conducted using STAR 2.5.4b ^86^ with default parameters, and the results were converted into the BAM format using Samtools 1.9 ^87^. Differential expression analysis was then carried out using the FeatureCounts count matrix ^88^, followed by DESEQ2 analysis ^89^, which utilized default parameters and prefiltering for comparison across samples. The total RNA and ChIPed samples were prepared in parallel.

### ChIP sequencing preparation

To ensure accurate normalization of the ChIP-seq data for H3K27me3, we incorporated a Spike-in technique. This involved adding a small amount of external chromatin to the experimental samples before conducting the ChIP reaction, which allowed us to standardize the signal from the experimental samples to the control signal in the final sequencing data ^90^. We did not use Spike-in technique for H3K9me3 because the abundance of H3K9me3 in its target regions may be relatively consistent, making it less susceptible to library preparation and sequencing depth variations, reducing the need for spike-in normalization.

The cells were grown to 70-80% confluency before being fixed in 4% formaldehyde for 10 min and stored at -80 °C. The resulting pellets were resuspended in 1 ml of ChIP-buffer, which contained 0.25% NP-40, 0.25% Triton X100, 0.25% Sodium Deoxycholate, 0.005% SDS, 50 nM Tris (pH 8), 100 mM NaCl, 5 mM EDTA, 1X PMSF, 2 mM NaF, and 1X cOmplete protease Inhibitor. The samples were then sonicated using a probe sonicator (Fisher Scientific Sonic Dismembrator Model 500) with 5 cycles at 20% power, 5 cycles at 25% power, and 5 cycles at 30% power. Each cycle lasted 10 sec, and the samples were kept on ice between each cycle to prevent overheating. The resulting lysates were then spun at high speed in a microcentrifuge for 30 min, and the protein concentration was measured using the BCA assay. The samples were diluted to a protein concentration of 2 mg/ml in ChIP-buffer. Then, 50 µl/ml of Protein G Plus-Agarose Suspension Beads were added to the samples for a 3-h incubation period to preclear them. 2% of the sample was collected as input and stored at -20 °C until DNA purification. For H3K27me3 ChIP, the remaining mixture was supplemented with 20 ng of spike-in chromatin and 2 ug of spike-in antibody. The mixture was then subjected to addition of the specific target antibodies and washed beads. Immunoprecipitation was carried out overnight at 4 °C with the mixture of samples, beads, and primary antibody (see Key resources table).

The resulting beads were washed once with Wash1, Wash2, and Wash3 buffers [0.10% SDS, 1% Triton X-100, 2 mM EDTA, 20 mM Tris (pH 8), with 150/200/500 mM NaCl for Wash1, Wash2, Wash3 buffer respectively], followed by a wash with Wash LiCl buffer [0.25 M LiCl, 1% NP-40, 1% Sodium Deoxycholate, 1 mM EDTA, 10 mM Tris (pH 8)], and two washes with TE buffer [10 mM Tris (pH 8), 1 mM EDTA]. The beads were then resuspended in elution buffer [1% SDS, 0.1 M NaHCO3], and the samples were de-crosslinked overnight at 65 °C. 20 µg of Proteinase K was added for 1 hour at 42 °C. DNA was then purified using a BioBasic DNA collection column, and DNA concentration was assessed via the Picogreen assay (Invitrogen). ChIPed DNA was sent to SickKids Genome centre for sequencing. The total RNA and ChIPed samples were prepared in parallel.

### ChIP sequencing processing

Quality control of reads and sequencing was assessed by FastQC (Babraham Bioinformatics). Adapters and low-quality reads were removed by Trimmomatics using the following parameters: ILLUMINACLIP:$Adapters:2:30:10 SLIDINGWINDOW:4:30 LEADING:30 TRAILING:30.

Alignment on hg19 human genome was performed using bowtie2 using “-end-to-end --phred33” parameters, and reads were directly sorted and converted to bam format by samtools. As sequencing was performed on different lanes to increase the depth, related reads on separate lanes were merged by samtools, and the quality of mapping was checked by samstat. Multiple filtering steps – including removal of low-quality aligned reads and reads aligned to mitochondria chromosome or any chromosome other than chr1-22, X, Y – were performed. Multi-mapped reads were also excluded.

As the broad signal of H3K27me3 is inaccurately represented by most peakcallers, we defined our peaksets by binning the genome at 1kb resolution and thresholding the top bins, as explained in ^83^, with an normalized read density above or equal to 100. Gained or lost signal were measured as the Log2FoldChange of the ratio between the average of the triplicate in each sample, two-tailed t-tests were used to establish significance. Genome binning and BigWig files used for visualization were generated by deepTools, bamCoverage function, excluded blacklisted regions and normalized using RPKM. For Spiked-In samples, the scaleFactor option, used the spike-in ratios, was used for each sample. Heatmaps, profile plot and tracks were generate using deepTools and samtools ^87,91^. Heatmaps and Profile plot were generated using 3 kb regions centered around the differential peakset. Both the computeMatrix and plotHeatmaps were run with default parameter; yMax, zMax and colors were adjusted in each condition to better represent the results.

To visualize ChIP-seq and RNA-seq dot plots and their correlation, dot plots were made by combining the RNA-Seq Log_2_FC between stimul. (veh) and mTOR-inhibited conditions and the Log_2_FC of the normalized binned reads density of any called peak annotated on that gene (+/-3 kb) or at the promoter regions, using clusterProfiler. Pearson correlation on the dot plot were performed using the ChIP-Seq Log_2_FC and RNA-Seq Log_2_FC of every peak colocalization with a gene.

### Cellular fractionation

Cytosolic, soluble nuclear and insoluble nuclear proteins were isolated using a commercial kit (#ab219177, Abcam) following the manufacturer’s instructions. Briefly, cells were washed twice in ice-cold phosphate-buffered saline (PBS) and lysed with cytosolic extraction buffer. After incubation on ice, nuclei were pelleted by centrifugation at 13,000 g for 1 min at 4 °C, and cytosolic proteins transferred to a clean microfuge tube. Nuclei were lysed using soluble nuclear lysis buffer, sonicated, and incubated on ice for 15 min, vortexing every 5 min. Insoluble material was pelleted by centrifugation at 13,000 g for 10 min at 4 °C, and the soluble nuclear fraction transferred to a clean microfuge tube. Isolated protein fractions were stored at −20 °C. Each buffer was supplemented with 100× protease inhibitor cocktail and DTT supplied with the kit.

### Flow cytometry for cell cycle and H3K27me3 analysis

The cells were harvested by trypsinization and subsequently washed twice with PBS. To render cells permeable, 70% ethanol was added dropwise while vortexing, and then the cells were stored at -20 °C until needed. After that, the cells were centrifuged at 900 g, washed once with cold PBS, and pelleted at 500 g. Next, the cells were washed with PBSA-T (5% BSA/0.1% Triton-100/PBS) at room temperature. The cells were then incubated with H3K27me3 antibody in PBSA-T for 1 hour at room temperature, followed by washing with PBS and incubation with secondary anti-Rat Goat Alexa Fluor 488 and 2 μg/ml DAPI analysis in PBSA-T for 30 min before analysis. A BD Fortessa (BD Biosciences) was used to collect a minimum of 10,000 events. All the experiments were carried out in at least three independent replicates. Key resources table outlines the reagents and antibodies that were used.

### Immunofluorescence (IF)

MCF7 cells were plated onto #1.5 coverslips of 12–15mm diameter in 12 well plates. Cells were allowed to recover overnight, followed by selected treatments. Coverslips were then washed with PBS twice, fixed with 2% paraformaldehyde/PBS for 20 min at room temperature, washed twice with PBS, and then followed by a 20 min fixation in 0.3% Triton-100/PBS solution. The coverslips underwent two washes with PBS and were then blocked with PBSA-T at room temperature for 1 hour. H3K9me3 antibody (1:1000 dilution) was then incubated for 1 hour at room temperature or overnight at 4 °C. Coverslips were washed twice with PBS and incubated with the fluorescent secondary antibody (1:750 dilution) with 2 ug/ml DAPI in PBSA-T for 1 hour at room temperature. Coverslips were washed twice in PBS, once in ddH2O and then mounted with Fluoromount-G (Invitrogen). Images were acquired with an LSM800 confocal microscope (Carl Zeiss AG) and analyzed as previously described ^92^. In brief, we measured the mean fluorescence intensity (MFI) of H3K9me3 foci relative to the background signal of the nucleus. Each data point represents a normalized signal on a per cell basis. Images were analyzed using the open-source Java ImageJ/Fiji program. Nuclei were first identified, by thresholding on DAPI fluorescence followed by analyzing particles larger than 75 µm^2^. H3K9me3 foci were counted using the Find Maxima feature and Measure function. At least 100 cells were analyzed per experiment. Key resources table outlines the reagents and antibodies that were employed, respectively. These experiments were performed in independent duplicate.

### Immunoprecipitation (IP)

The cells were grown in a 15cm cell culture dish and washed twice with ice-cold PBS before being scraped. The cells were then lysed using lysis buffer containing 50 mM Tris-HCl pH 7.5, 150 mM NaCl, 0.5% NP-40, 2 mM EDTA, 0.5 mM DTT, 1 mM PMSF and 1× EDTA-free cOmplete protease inhibitor cocktail (Roche Applied Science). The protein concentration was determined via BCA assay, and 1 mg of protein was diluted in 1 ml of lysis buffer. Protein G agarose beads were washed with lysis buffer. To pre-clear the lysates, they were incubated with protein G agarose beads, followed by centrifugation to pellet the beads. 10 % of the sample is collected as input and kept at -20 °C. Immunoprecipitation was performed using anti-EZH2 rabbit polyclonal antibody or rabbit IgG as a control, according to the manufacturer’s protocol, with 1 hour incubation. The washed beads were then added to the tube and incubated for 1 hour with rotation. The mixture was washed three times with the lysis buffer, and the beads were resuspended in 2X SDS loading buffer. Total cell lysates and immunoprecipitates were separated by SDS-PAGE and analyzed by Western blotting. Key resources table outlines the reagents and antibodies that were employed, respectively. These experiments were performed in independent triplicate.

### Quantification and statistical analysis

For statistical analysis one-way ANOVA (Prism; Graphpad) was employed, unless indicated otherwise. Before statistical analysis, technical replicates were averaged. Statistical comparisons were made by analyzing 2-3 independent experiments, each incorporating the average of technical replicate outcomes. Data was presented as mean ± SD from independent experiments. Further specifics concerning data quantification, presentation, and method-specific statistical analysis can be found in the respective STAR Methods sections and figure legends.

## KEY RESOURCES TABLE

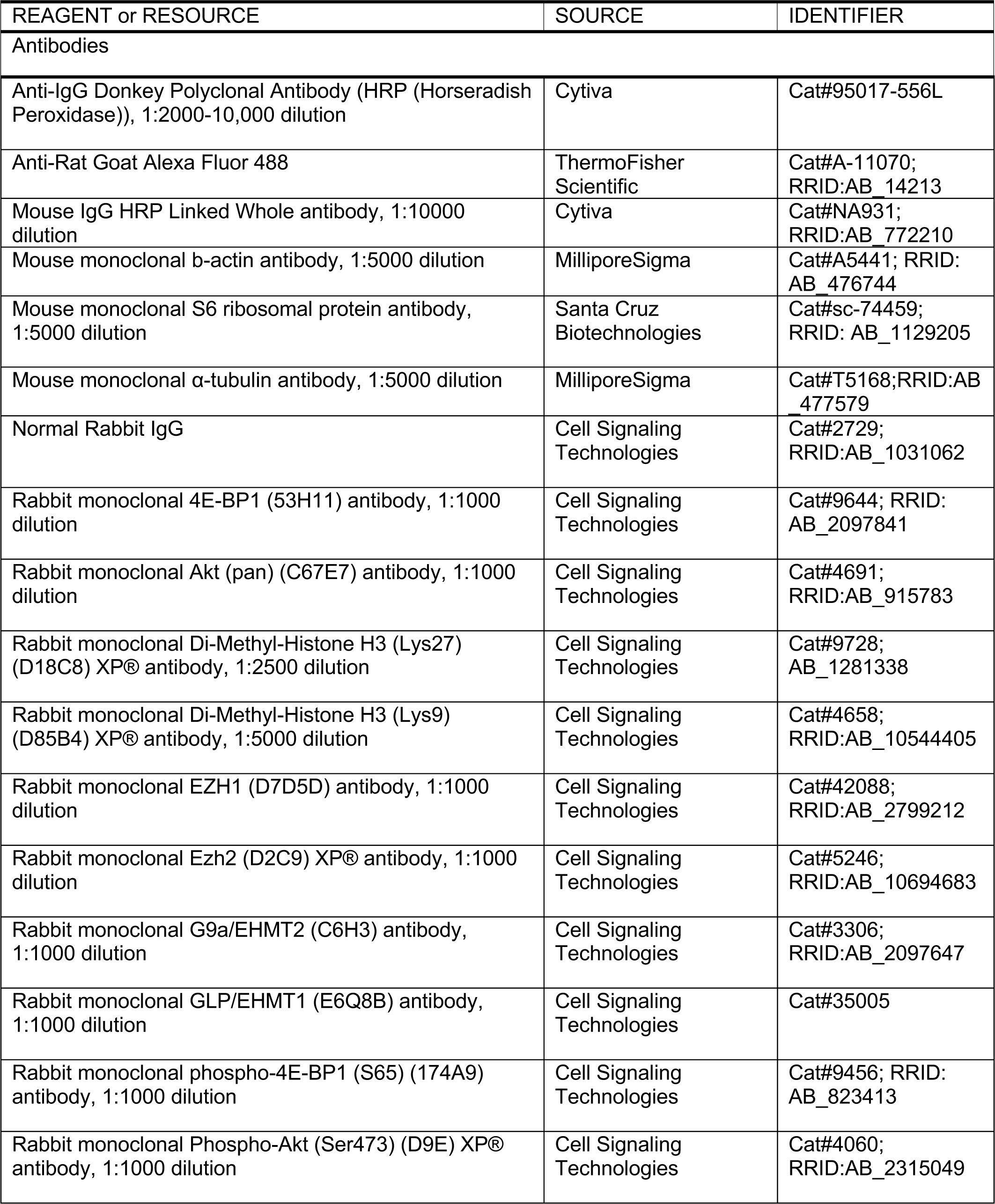
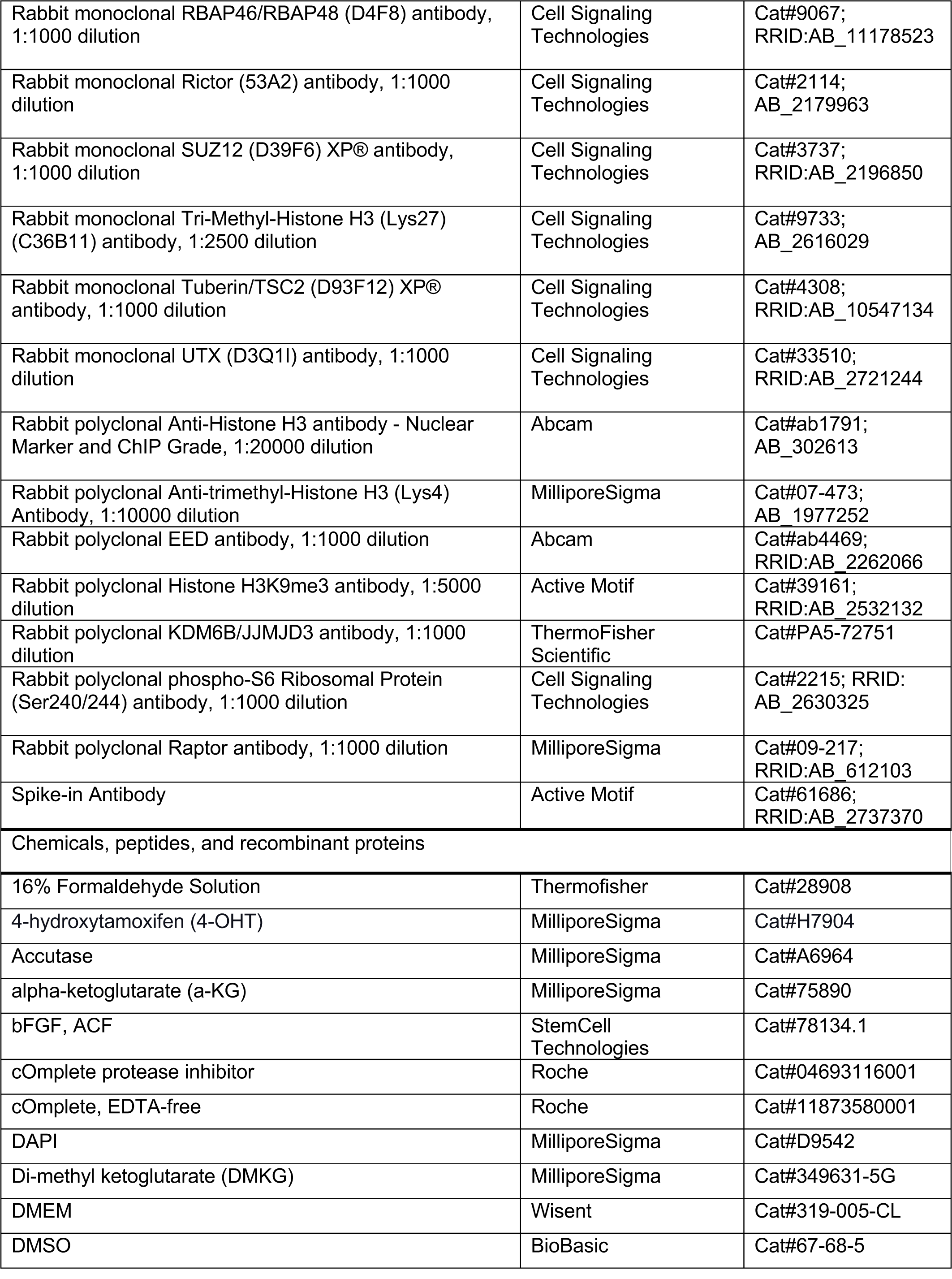
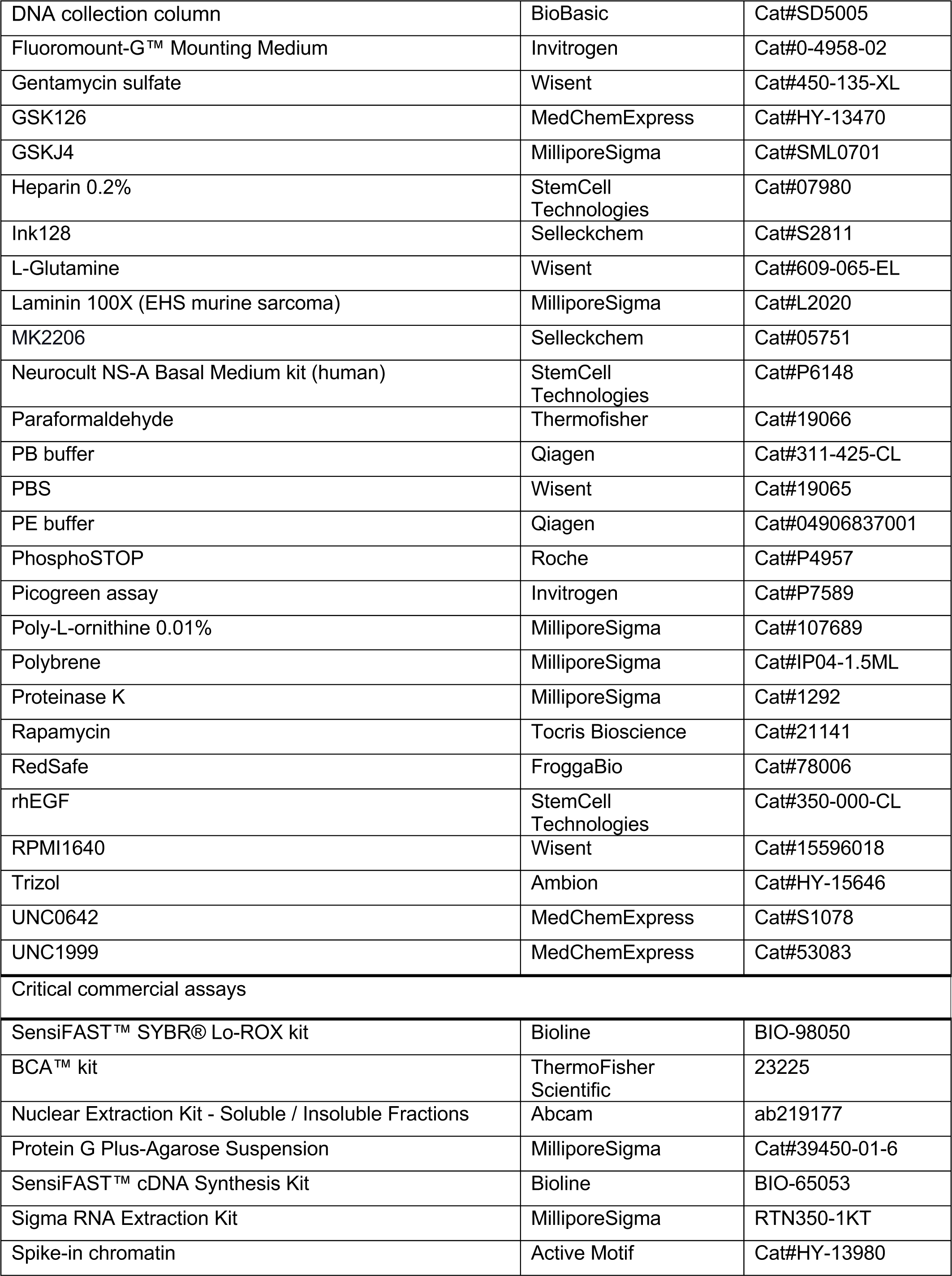
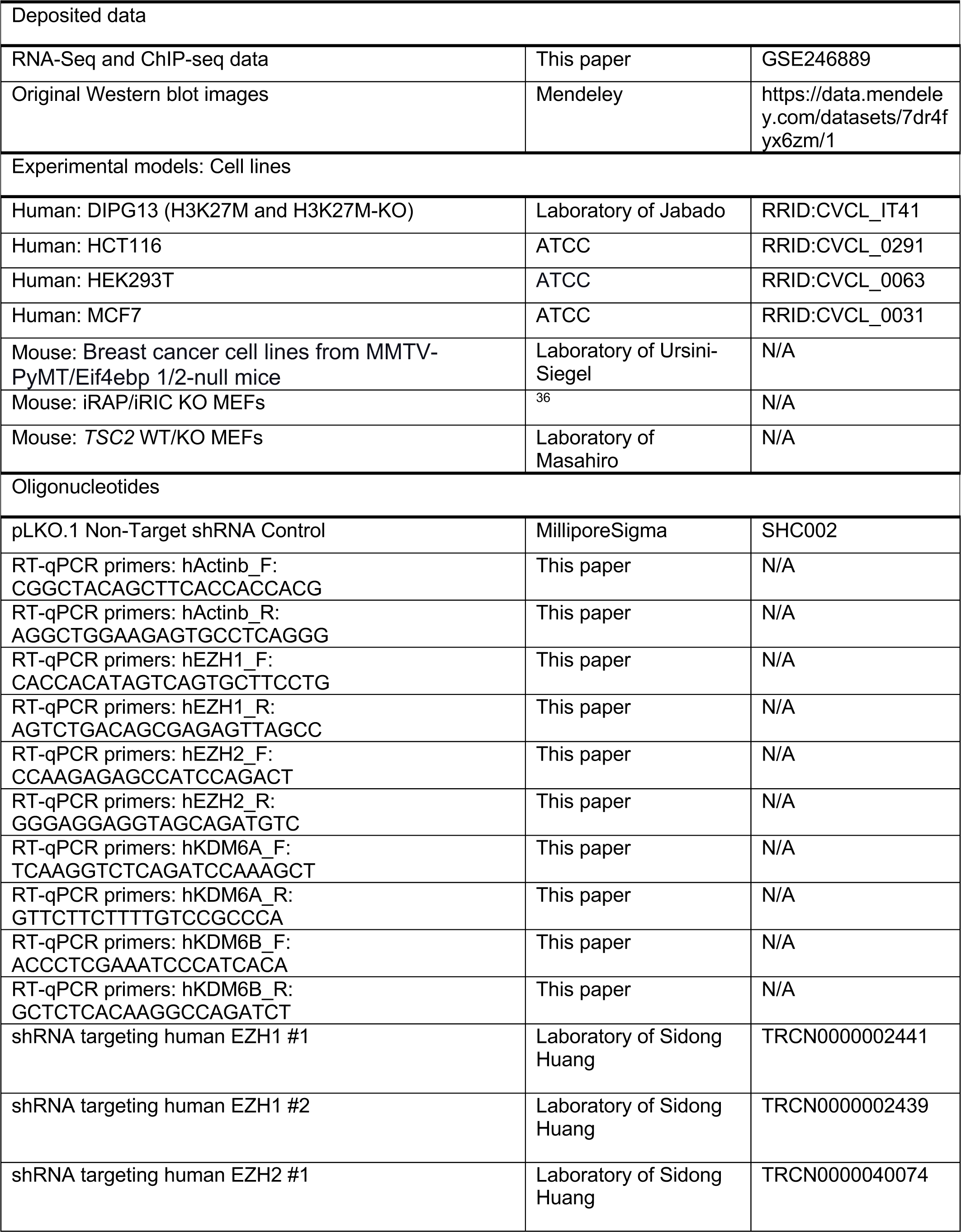
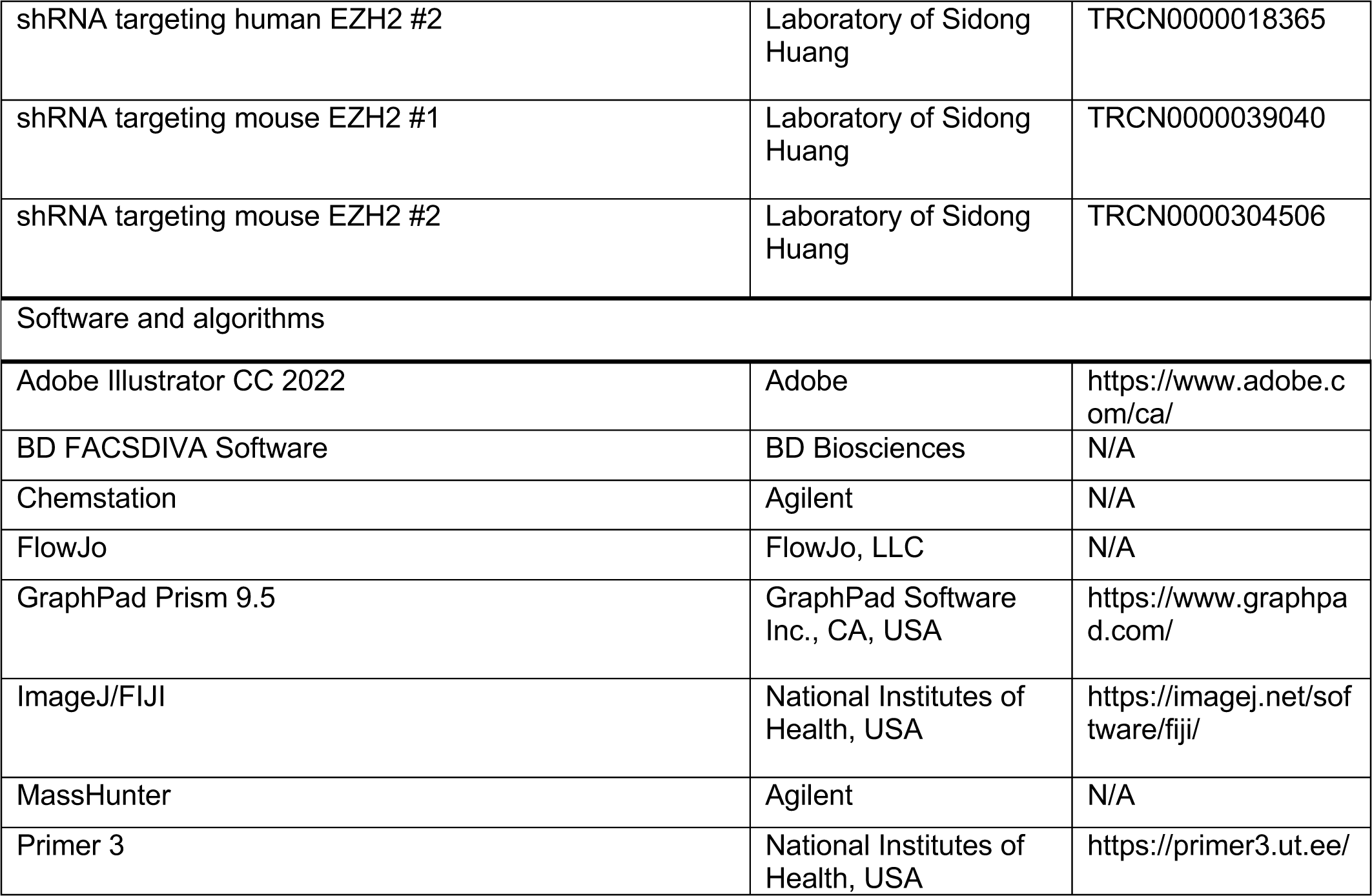

