## Supplementary Figures for "MTOR modulation induces selective perturbations in histone methylation which influence the anti-proliferative effects of mTOR inhibitors"

### Supplementary Data

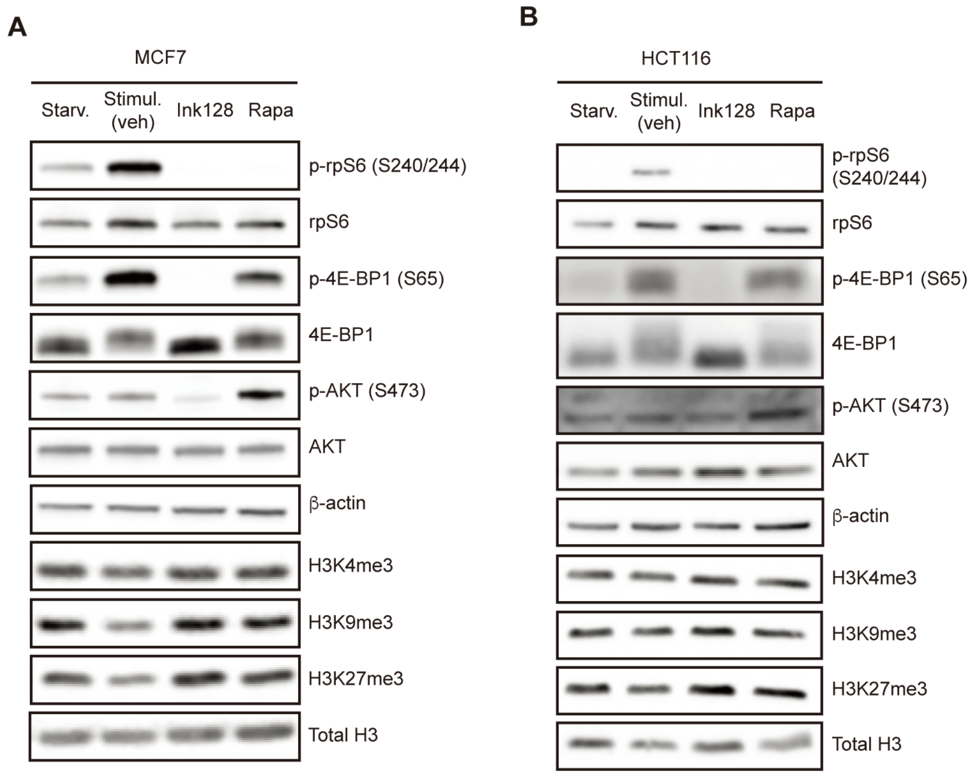


**Figure S1. The effect of serum starvation in MCF7 and HCT116.**

(A-B) MCF7 (A) and HCT116 (B) cells were serum-starved overnight and then stimulated with 10% FBS in the presence of Ink128 (100 nM) or rapamycin (50 nM) for 48 hours. Levels of the indicated proteins were monitored by Western blotting (n=3). β-actin served as a loading control.

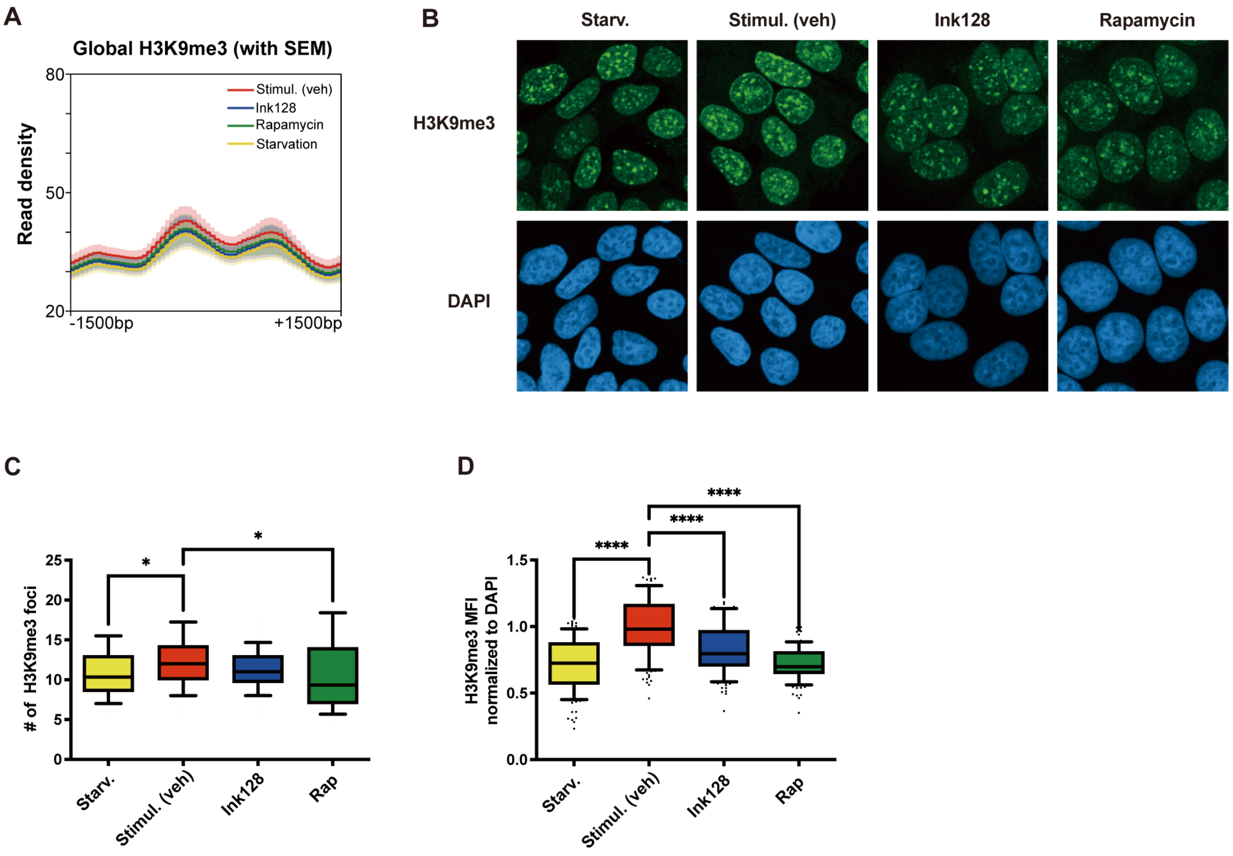


**Figure S2. The effects of mTOR inhibition on induction of H3K9me3 are modest**

(A) The normalized average read peak density profiles for H3K9me3 with SEM in MCF7 treated as indicated. (B) Representative confocal micrograph images of IF staining for H3K9me3 (green) and DAPI (blue) in MCF7 cells that were serum-starved overnight and then simultaneously treated with a vehicle (DMSO) or Ink128 (100 nM) in the presence of 10% FBS, or were kept serum-starved for 48 hours. Cells were fixed, stained, and imaged via confocal microscopy. (C-D) Quantification of H3K9me3 foci (C) and the mean fluorescence intensity (MFI) (D). At least 100 cells were analyzed per experiment. Bars represent mean ± SD. *$P < 0.05$, ****$P < 0.0001$, one-way ANOVA with Dunnett's post-test. All IF staining was carried out in three independent replicates (n=3).

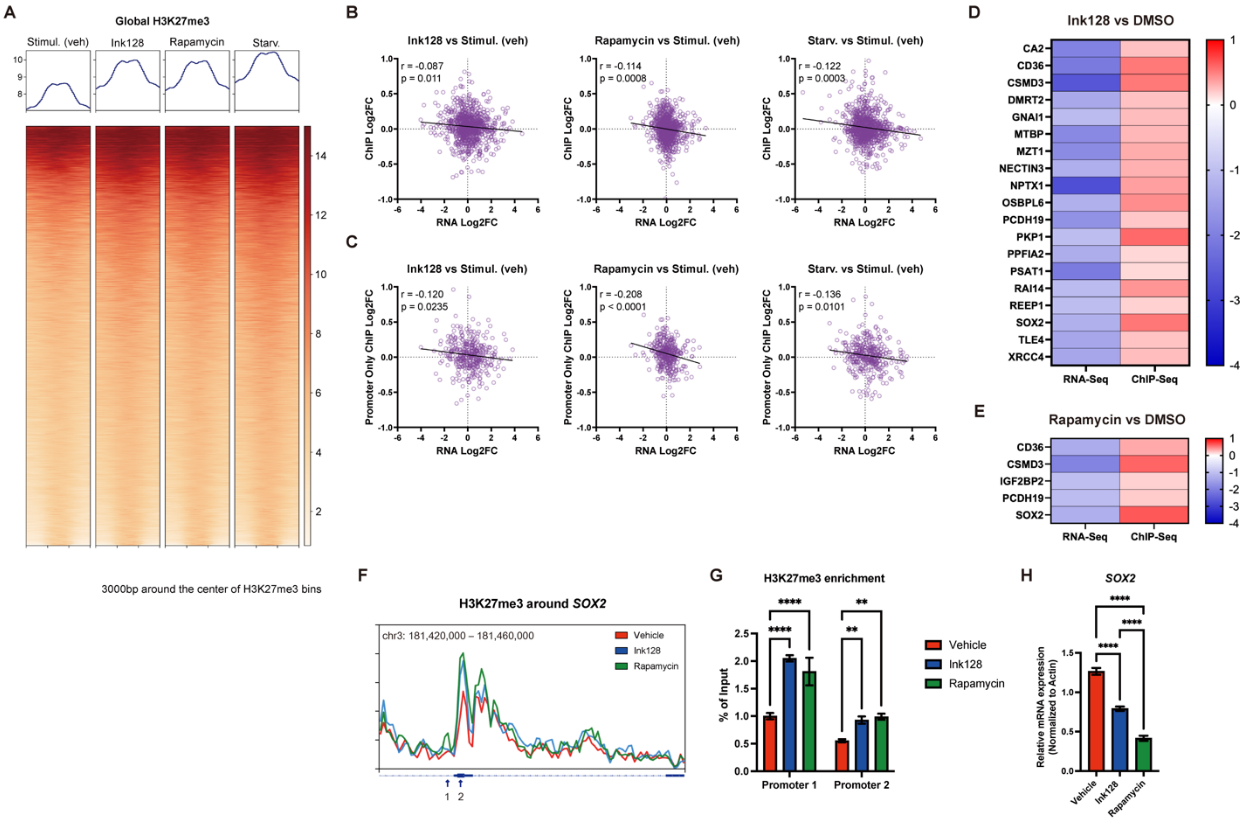


**Figure S3. The ChIP-seq and RNA-seq analysis of the effects of mTOR inhibition in MCF7 cells.**

(A) Heatmap plots of ChIP-seq signal intensity for H3K27me3 in MCF7 treated as indicated. (B-C) Correlation between ChIP-seq and RNA-seq data across all regions (B) and specifically in the promoter region (C). (D-E) Heatmap plots display genes that have gained H3K27me3 (within 3kb of the gene) and exhibit downregulated RNA-Seq results following treatment with Ink128 (D) or rapamycin (E) compared to the vehicle. RNA-Seq Log2FC threshold: -1, basemean threshold: 50, ChIP-Seq Log2FC threshold: 0.15 (F) H3K27me3 levels determined by ChIP-seq analysis at the SOX2 loci. (G) H3K27me3 ChIP performed prior to qPCR at promoter regions for the genes indicated. **P< 0.01, ****P< 0.0001, one-way ANOVA with Dunnett's post-test compared to vehicle. (H) MCF7 cells were serum-starved overnight and then stimulated with 10% FBS in the presence of Ink128, rapamycin or a vehicle (DMSO) for 48 h. *SOX2* mRNA levels were assessed by RT-qPCR (n=3). ****P< 0.0001, one-way ANOVA with Dunnett's post-test compared to vehicle.

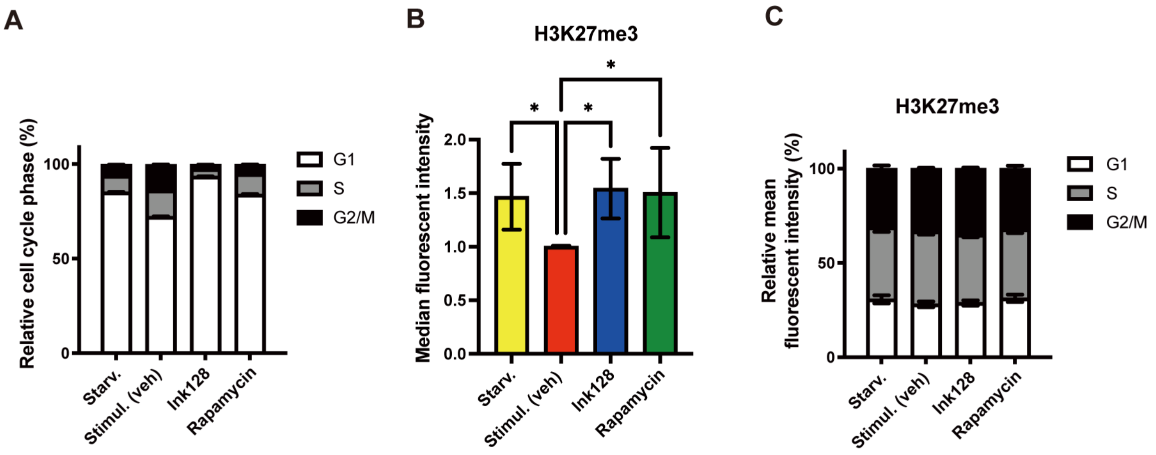


**Figure S4. The alteration of H3K27me3 is not mediated by the cell cycle changes following mTOR inhibition.**

(A) Cell cycle profiles of MCF7 cells that were serum-starved overnight (Starv-yellow) and then treated with a vehicle [DMSO(Stimul-red)], Ink128 (100 nM;blue) or rapamycin (50 nM;green) in the presence of 10% FBS were monitored by flow cytometry by using DAPI staining. (B) The overall mean fluorescent intensity of H3K27me3 in MCF7 cells treated as in panel (A) was assessed by flow cytometry. *P< 0.05, one-way ANOVA with Dunnett's post-test. (C) The mean fluorescent intensity of H3K27me3 in the indicated phases of cell cycle was determined by flow cytometry. Herein, MCF7 cells were treated as described in panel (A). Bars represent mean ± SD. All flow cytometry was carried out in three independent replicates (n=3).

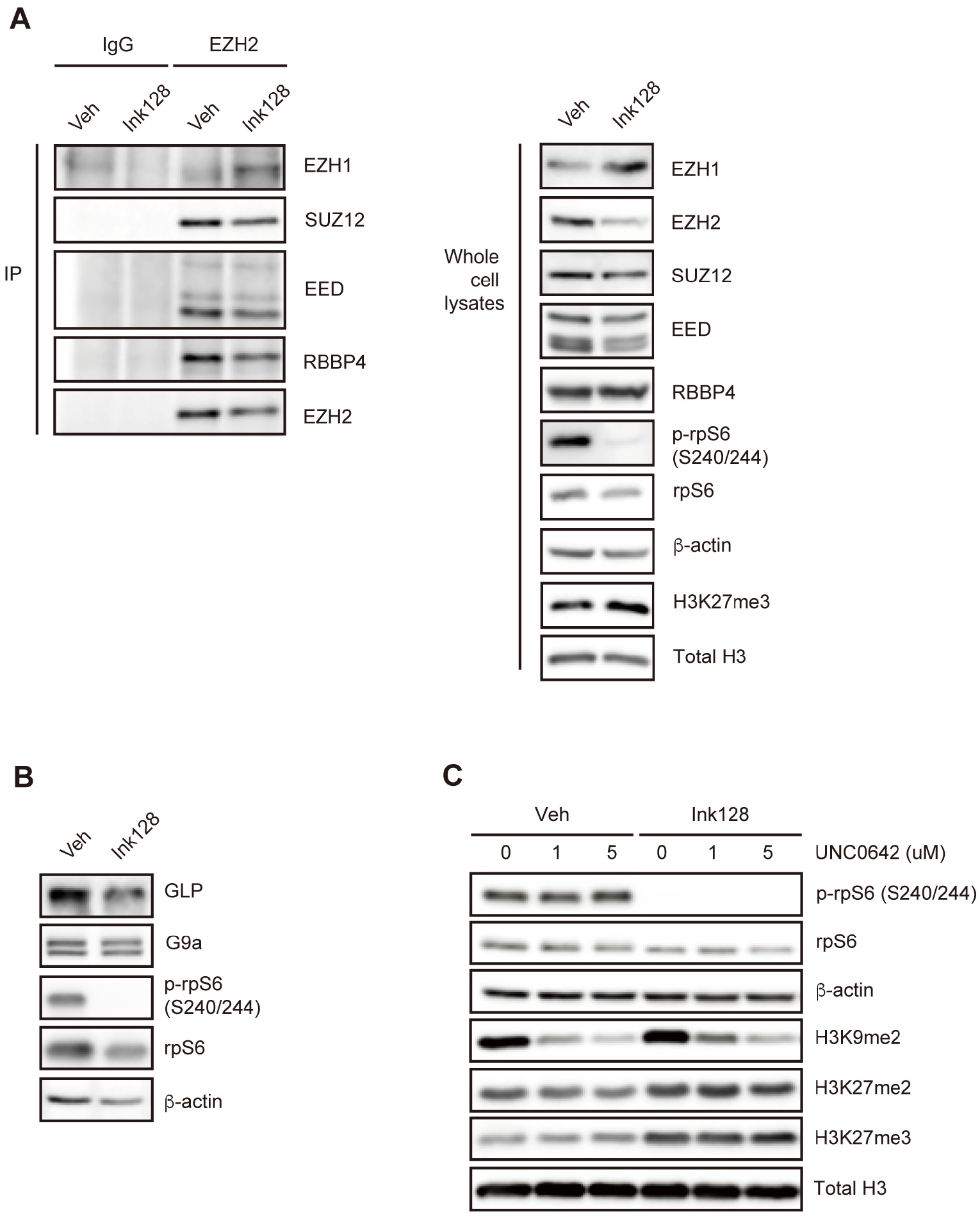


**Figure S5. Although PRC2 activity is required for H3K27me3 induction upon mTOR inhibition, mTOR inhibitors do not affect PRC2 assembly or GLP/G9a.**

(A) Immunoblot analysis of whole cell lysates and anti-EZH2 immunoprecipitates derived from MCF7 cells treated with 100 nM Ink128 for 24 hours after overnight serum-starvation. β-actin served as a loading control. (B) Levels of the indicated proteins in MCF7 that were serum-starved overnight and then treated with a vehicle (DMSO) or Ink128 (100 nM) in the presence of 10% FBS for 48 hours were determined by Western blotting (n=3). β-actin served as a loading control. (C) Levels of the indicated proteins in MCF7 treated with indicated concentrations of UNC0642 in the presence of 100 nM Ink128 or a vehicle (DMSO) for 48 hours were determined by Western blotting (n=2). β-actin served as a loading control.

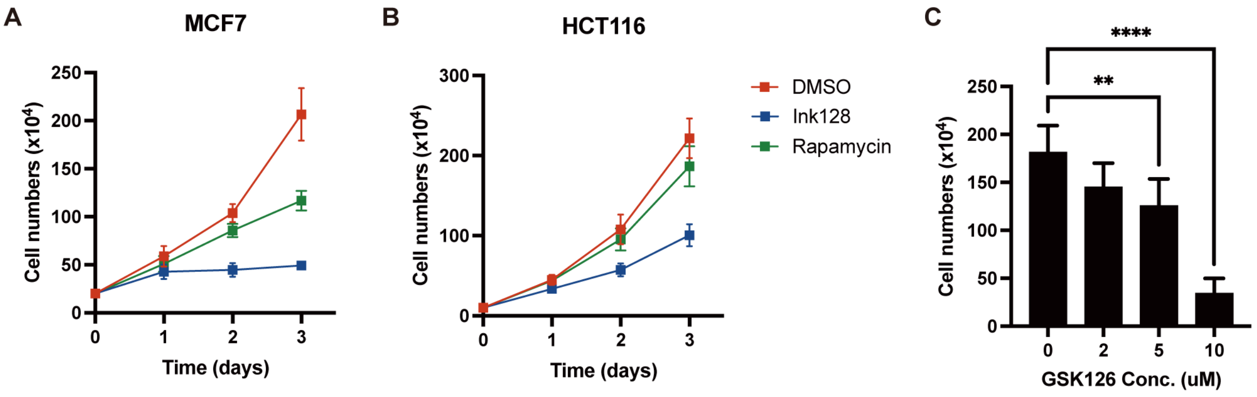


**Figure S6. Cell proliferation decreases upon mTOR inhibition or H3K27me3 reduction.**

(A-B) Proliferation curves of the indicated cancer cell lines treated with vehicle (DMSO), 100 nM Ink128 or 50 nM rapamycin for denoted time periods (n=3). Bars represent mean ± SD. (C) Counting of viable MCF7 cells treated with indicated concentrations of GSK126 for 72 hours (n=3). Bars represent mean ± SD. ****P< 0.0001, one-way ANOVA with Dunnett's post-test.
